# Phase diversity-based wavefront sensing for fluorescence microscopy

**DOI:** 10.1101/2023.12.19.572369

**Authors:** Courtney Johnson, Min Guo, Magdalena C. Schneider, Yijun Su, Satya Khuon, Nikolaj Reiser, Yicong Wu, Patrick La Riviere, Hari Shroff

## Abstract

Fluorescence microscopy is an invaluable tool in biology, yet its performance is compromised when the wavefront of light is distorted due to optical imperfections or the refractile nature of the sample. Such optical aberrations can dramatically lower the information content of images by degrading image contrast, resolution, and signal. Adaptive optics (AO) methods can sense and subsequently cancel the aberrated wavefront, but are too complex, inefficient, slow, or expensive for routine adoption by most labs. Here we introduce a rapid, sensitive, and robust wavefront sensing scheme based on phase diversity, a method successfully deployed in astronomy but underused in microscopy. Our method enables accurate wavefront sensing to less than λ/35 root mean square (RMS) error with few measurements, and AO with no additional hardware besides a corrective element. After validating the method with simulations, we demonstrate calibration of a deformable mirror > 100-fold faster than comparable methods (corresponding to wavefront sensing on the ~100 ms scale), and sensing and subsequent correction of severe aberrations (RMS wavefront distortion exceeding λ/2), restoring diffraction-limited imaging on extended biological samples.

## Introduction

Fluorescence microscopy is invaluable in biological research due to its contrast, resolution, speed, and potential for live imaging. However, the refractile nature of biological tissues or misaligned optics often cause undesirable bending of illumination and emission light, introducing wavefront distortion (**Fig. 1a**). Such ‘optical aberrations’ prevent diffraction-limited imaging, degrading image contrast, resolution, and signal.

**Fig. 1.**
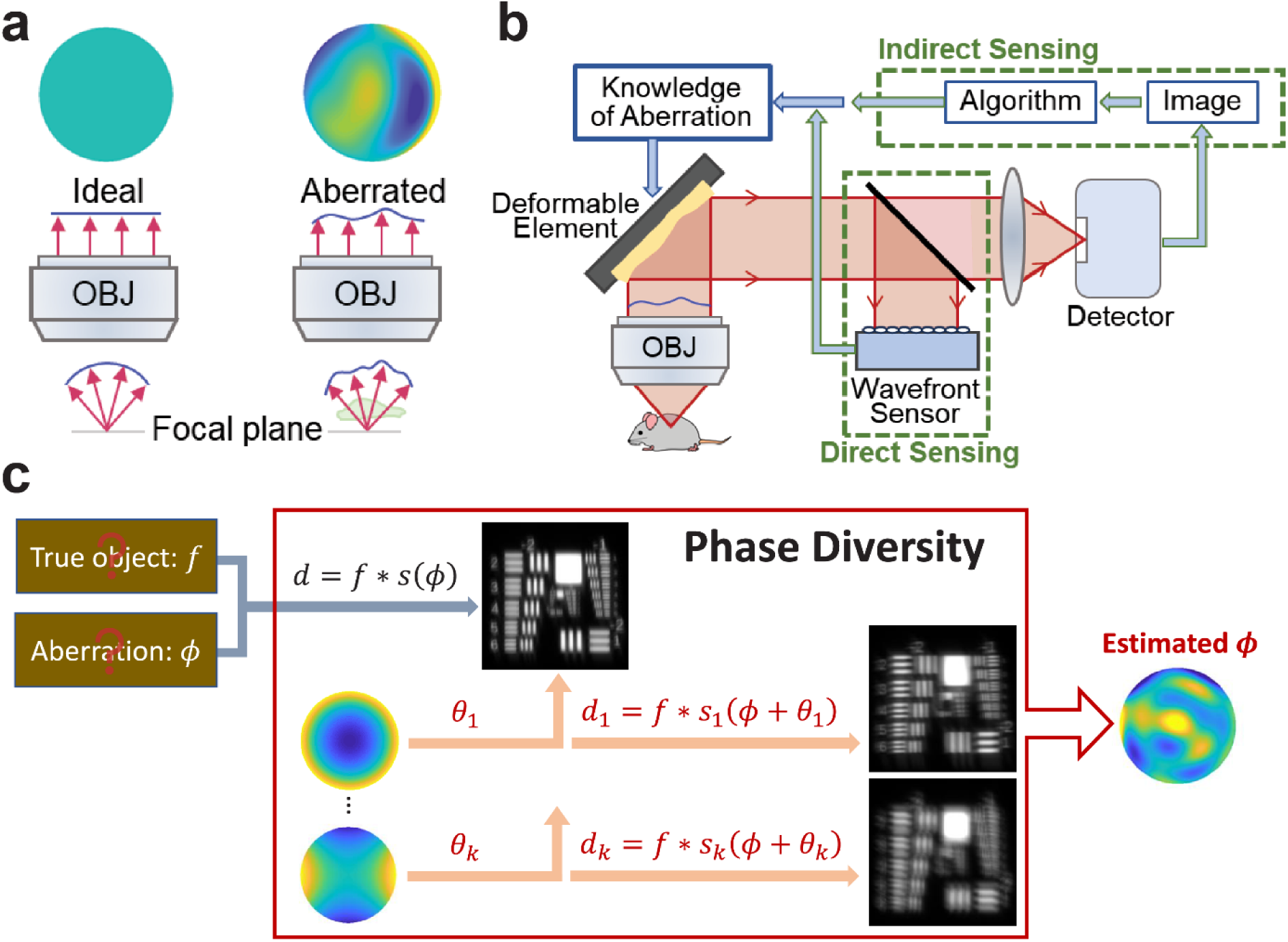
Sensing aberrated wavefronts in fluorescence microscopy. **a)** Left: In an ideal fluorescent sample, spherical wavefronts are captured from the sample and converted to parallel wavefronts by the objective lens (OBJ), yielding a flat wavefront at the pupil plane (top left). Right: in most real refractile samples, bending of light yields a distorted wavefront with noticeable phase variation at the pupil (top right). **b)** Adaptive optics techniques, which sense the aberrated wavefront and use this knowledge to apply an equal and opposite corrective wavefront using a deformable element, can be broadly divided into two classes based on the method of wavefront sensing. In direct sensing (bottom), a dedicated wavefront sensor is used in a separate optical path to sense aberrations. By contrast, indirect sensing (top) uses the same optical path as for fluorescence imaging, employing algorithms to sense the wavefront based on fluorescent images of the sample. **c)** In this work, we develop a phase diversity method for wavefront sensing in fluorescence microscopy. Given the unknown object *f* and unknown aberrated point spread function *s(ϕ)*, the image *d* does not contain sufficient information to infer the unknown aberration. In phase diversity, additional aberrated images *d*_*1*_,…,*d*_*k*_ are collected, each with known diversity aberrations *θ*_1_,…,*θ*_*k*_ purposefully added. The set of diversity images *d*_*1*_,…,*d*_*k*_ now contribute additional information, enabling us to estimate the unknown aberration *ϕ*.

Adaptive optics technologies (AO^1,2^) attempt to restore imaging performance by (i) sensing the distorted wavefront and (ii) applying a corrective wavefront of equal amplitude but opposite sign (typically via an adaptive element such as a deformable mirror or spatial light modulator), thereby suppressing aberrations. The key to implementing effective AO is the sensing, which would ideally be performed as rapidly and accurately as possible. Despite its importance, there is no consensus on best practices for wavefront sensing, which remains an active area of research.

Wavefront sensing methods can be broadly classified as ‘direct’ or ‘indirect’, which offer their own advantages and disadvantages (**Fig. 1b**). Both classes have been used effectively for AO in confocal^3-5^, multiphoton^6-10^, light sheet^11-13^, and super-resolution imaging^14-16^. Direct approaches use dedicated hardware (often a Shack-Hartmann wavefront sensor, SHWFS) to rapidly sense the wavefront, and are most effective in conjunction with point-like sources (‘guide stars’) that are usually exogenously introduced^6,17,18^. Typical disadvantages of direct approaches include the need to form an image (which requires sufficient ballistic signal to reach the detector), limiting these methods to weakly scattering samples; the cost of the sensing hardware (which can be considerable^5^); low sensitivity of the sensing hardware; and differential aberrations in the sensing vs. imaging paths.

By contrast, indirect sensing methods reconstruct the wavefront from a sequence of intensities or images, each with a corresponding change applied to the adaptive element^3,8-10,19^. Advantages of indirect approaches include their relatively low cost and simple hardware (since no wavefront sensor is needed) and improved performance in opaque tissue (since the requirement for ballistic signal is usually relaxed). The Achilles heel of indirect methods is their slow speed, as the need for many sequential measurements and the associated computational burden results in slow wavefront sensing that often far exceeds the time for image acquisition.

Within the plethora of indirect sensing methods, phase diversity methods (PD^20-23^) are under-explored in fluorescence microscopy. Originally developed for astronomical applications, phase diversity approaches can efficiently estimate both an extended object and the associated aberrated wavefront from as few as two images. Despite its potential, we are aware of only a handful of studies that have attempted to apply PD-based sensing to biological samples^24-26^.

Here we present a rapid and accurate implementation of PD and demonstrate its use in wavefront sensing and AO. First, we conducted simulations to benchmark our algorithm, exploring the regime in which the algorithm excels. Second, we built a simple widefield microscope incorporating a deformable mirror, demonstrating that our PD implementation can calibrate the mirror orders of magnitude faster than comparable indirect methods, and faster than a commercially available Shack-Hartmann wavefront sensor (SHWFS) while providing similar accuracy (producing a flat wavefront within 15 nm root mean square (RMS) error). Finally, we show that PD can be used in an AO loop, correcting severe optical aberrations induced within biological specimens, thereby restoring diffraction-limited imaging.

## Results

### Phase diversity for fluorescence microscopy

In fluorescence imaging, we can represent the formation of the image *d* as a convolution between the object *f* and the point spread function *s(ϕ), d* = *f* * *s(ϕ)*. Here *ϕ* represents a phase aberration, we have neglected noise, and have assumed that *s* is spatially invariant. For AO applications it would be desirable to estimate *ϕ* to compensate for it, thereby improving image quality. In general, *f* and *ϕ* are both unknown and thus we cannot estimate either from the single image *d*. Phase diversity methods address this problem with the collection of additional diversity images *d*_*1*_,*…,d*_*k*_ in which known diversity aberrations *θ*_*1*_,*…,θ*_*k*_ are purposefully introduced. The additional information contained within such images allows us to estimate *f* and the unknown aberration *ϕ* (**Fig. 1c**).

Previous applications of phase diversity have mainly focused on astronomical applications in which aberrations are caused by atmospheric turbulence. We developed an implementation well suited to fluorescence microscopy (**Supplementary Note 1**). Given the original aberrated image *d*, the ideal PSF *s*, the additional diversity images *d*_*1*_,*…,d*_*k*_ and the known diversity aberrations *ϕ*_*1*_,*…,ϕ*_*k*_, our algorithm uses the Gauss-Newton algorithm to rapidly estimate both *f* and *ϕ* (**Methods, Supplementary Fig. 1**).

We used simulations to initially benchmark our algorithm, and to characterize its performance (**Fig. 2, Supplementary Figs. 2, 3**). We added synthetic aberrations and noise to a variety of noiseless phantom objects to produce aberrated images, finding that using a single additional diversity image was sufficient to estimate the aberration, the underlying object, and to enable image correction via AO (**Fig. 2a-c, Supplementary Fig. 2a**). Simulations also allowed us to investigate the relative efficacy of different types of diversity aberrations (**Fig. 2d, Supplementary Fig. 3**), the effect of increasing the number of diversity images (**Fig. 2e**), and the influence of signal-to-noise ratio (SNR) in the aberrated image (**Supplementary Fig. 2c**). In addition, simulations aided us in selecting model parameters for our algorithm (**Methods, Supplementary Fig. 2b**).

**Fig. 2.**
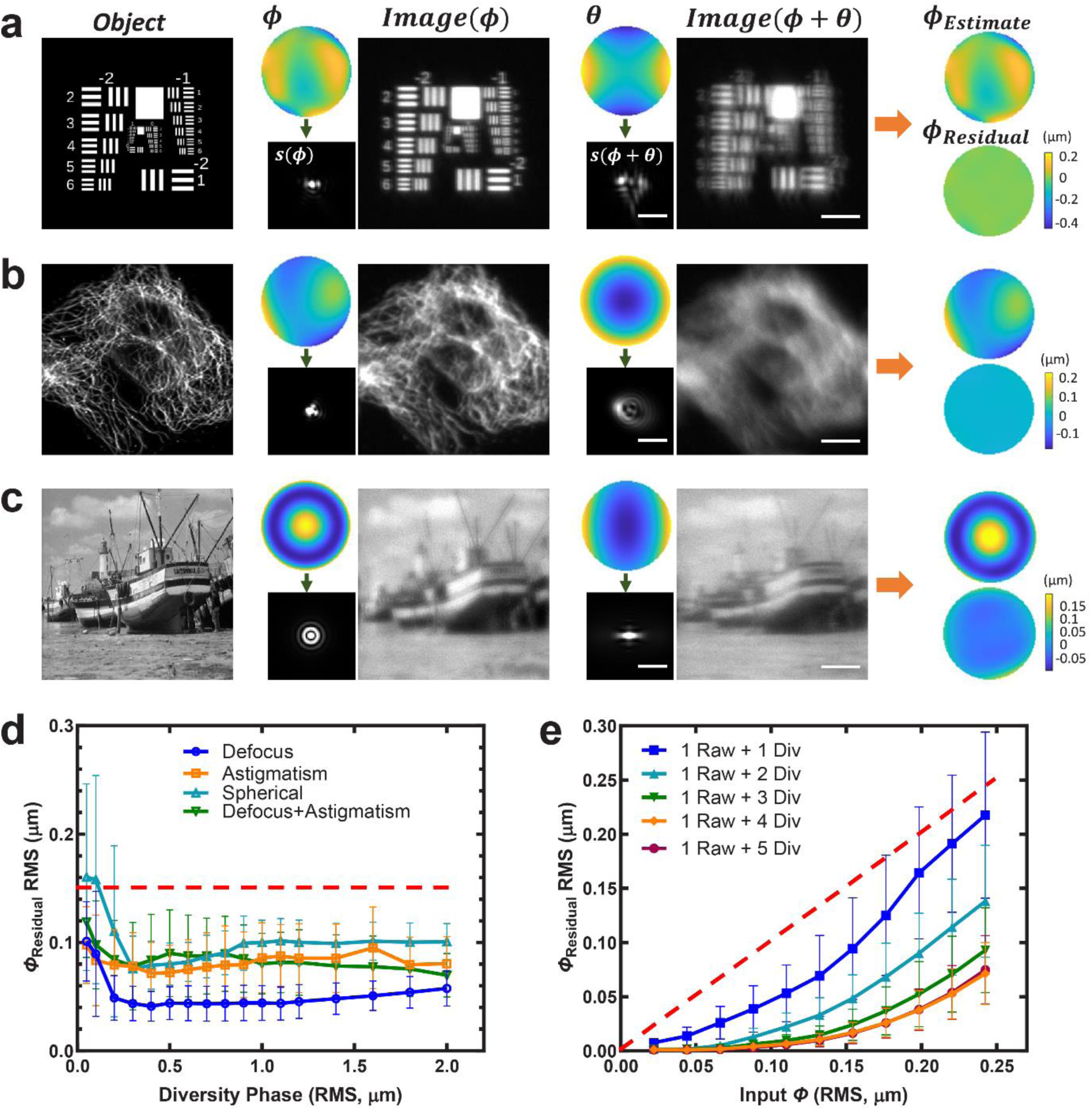
Simulations confirm that phase diversity enables accurate wavefront sensing. Different objects (1951 USAF resolution test chart, immunolabeled microtubules in fixed cell, Fishing Boat, **a-c)** respectively, left column) are contaminated with aberrations (wavefront *ϕ* and corresponding point spread function (PSF) *s(ϕ)*) and noise, yielding aberrated, noisy images (*Image(ϕ)*). Acquiring additional diversity images (*Image(ϕ+θ)*) with additional known diversity phases *θ* enables accurate phase diversity-based estimation of the original aberration (*ϕ*_*Estimate*_) with small residual wavefront error (*ϕ*_*Residual*_). **d)** Residual root mean square (RMS) wavefront error for different choices of diversity phase, as a function of the magnitude and type of the diversity phase. Here the input RMS wavefront distortion was 0.15 μm (dashed red line), and the signal-to-noise ratio (SNR) of the aberrated image was 10. See also **Methods, Supplementary Fig. 3**. Means and standard deviations from 50 simulations are shown. **e)** Residual RMS wavefront error for different numbers of diversity images, as a function of the magnitude of input aberration. Dashed red line indicates the y = x line, i.e., images corresponding to points underneath this line have less wavefront distortion than the original image. SNR of the aberrated image is 10, see also **Supplementary Fig. 2**. Means and standard deviations from 50 simulations are shown. Scale bars in **a)** 5 μm for objects and images, 2 μm for PSFs. Results in the last column of **a-c)** and in **d)** are obtained by using two images in our phase diversity algorithm, the original aberrated image and one diversity image. Simulations in **d, e)** are based on the USAF test target shown in **a)**. See also **Methods**.

For example, we found that astigmatism, primary spherical, or defocus Zernike basis functions were effective as diversity aberrations (**Fig. 2d**), whereas coma and trefoil were not (**Supplementary Fig. 3**). Using more diversity images provided better aberration estimation than fewer (**Fig. 2e**), although for the diversity phases we chose, performance saturated at four diversities (for a total of five images, including the initial aberrated image). Finally, the quality of wavefront sensing deteriorated at lower input image SNR, although the algorithm provided usable wavefront estimation for input root-mean-square (RMS) wavefront distortions up to ~0.25 μm even with input image SNR as low as ~3 (**Supplementary Fig. 2c**).

### Rapid sensorless calibration of a deformable mirror

To test the performance of our method in an experimental setting, we built a widefield fluorescence microscope incorporating a widely used, commercially available deformable mirror (DM) in a conjugate pupil plane (**Supplementary Fig. 4**). The DM uses 52 electromagnetic actuators to manipulate the shape of a reflective membrane (**Fig. 3a**), thereby introducing or correcting aberrations, but relies on accurate calibration of each actuator before use. Calibration in this context means determining the ‘influence function’ of each actuator (the induced wavefront as a function of applied voltage). Once known, the influence functions can be inverted, thereby generating a ‘control matrix’ that provides the per-actuator voltage needed for a desired aberration (**Methods**, ref.^27^). Since calibration relies on accurate wavefront sensing, it provides an excellent initial test of our phase diversity method.

**Fig. 3.**
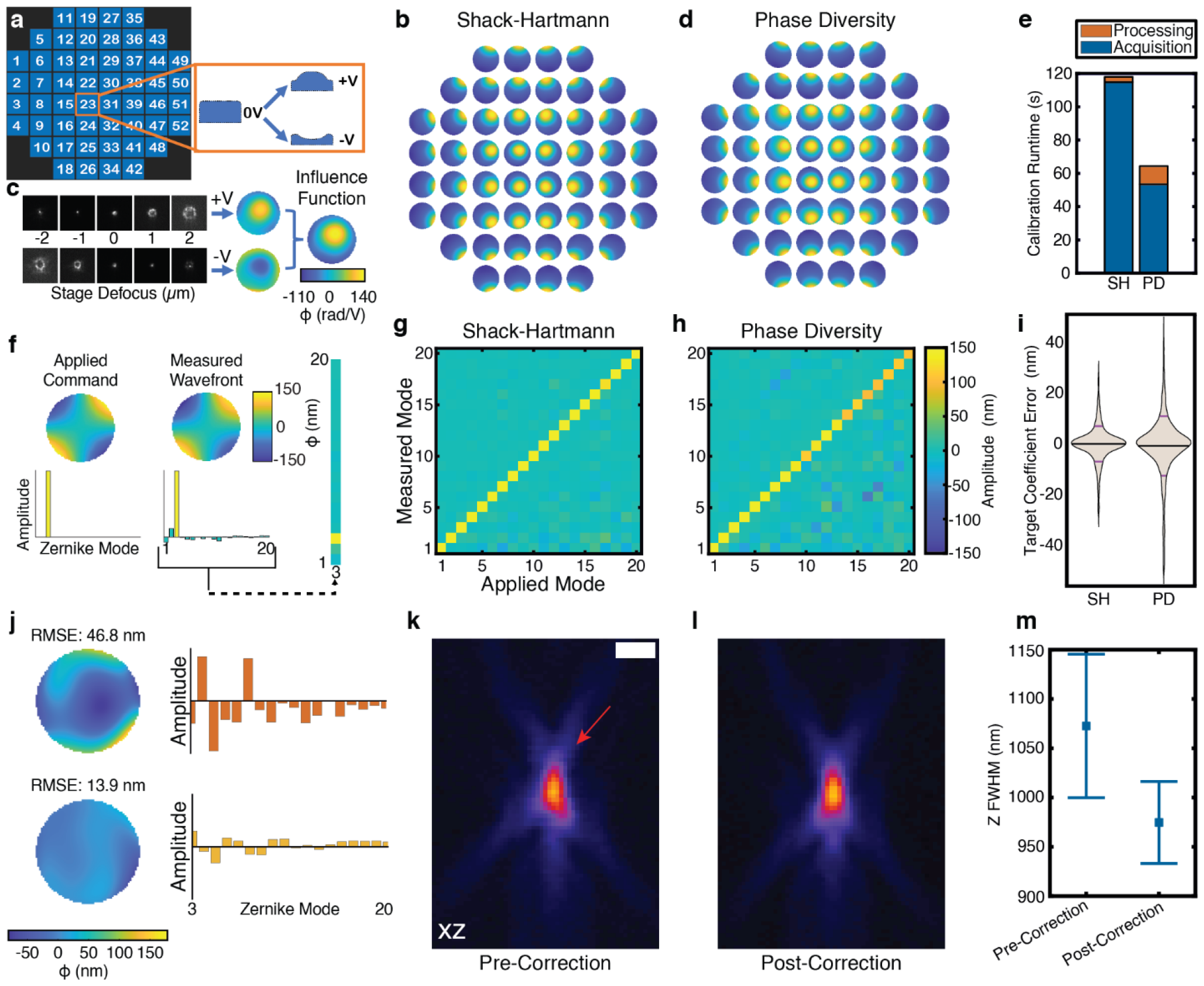
Phase diversity enables rapid calibration of a deformable mirror. **a)** Our 52-element deformable mirror (left) is calibrated by applying positive and negative voltages (±V) to each actuator (right, shown for actuator 23) and measuring the induced wavefront as a function of applied voltage (‘influence function’). **b)** Influence functions measured using direct wavefront sensing with a commercially available Shack-Hartmann wavefront sensor (SH). **c)** For calibration using phase diversity (PD), we acquire a series of widefield images of 500 nm beads at different axial positions (defocus diversities) of our stage (±2, ±1, 0 μm) for each voltage, and use the resulting wavefronts to compute influence functions **d)** that closely resemble those acquired with SH. **e)** Despite a larger computational burden, PD results in a faster calibration (64 s) than SH (118 s) due to the shorter acquisition time. **f)** To further investigate the accuracy of our calibrations, we command the deformable mirror to issue Zernike modes from 1-20 (left, shown for Zernike mode 3, oblique astigmatism), and then measure the resulting wavefront with the SH. Ideally, the measurement would show only the target Zernike mode (yellow), but in practice (middle) we also measure some degree of off-target Zernikes (green). Top row: wavefront representation; bottom row: amplitudes for each Zernike mode. The measurement of the response to each Zernike mode (1-20, ANSI convention) represents a single column in our characterization assay matrix (right). Characterization assays for SH- **g)** and PD- **h)** derived calibrations closely resemble each other, although the latter produces a noisier assay. Note that the SH was used to measure responses in both cases, see also **Supplementary Fig. 6e, f** for an example where PD is used. This is confirmed in quantitative assessment **i)** of the errors between target and measured coefficient values for each element in SH and PD characterization matrices. Violin plots are shown, with mean (horizontal black bars) and standard deviations (horizontal red bars) after outlier removal, SH: 0 ± 7 nm; PD: -1 ± 12 nm (see also **Supplementary Fig. 6**). The PD-derived calibration is of sufficient quality that it can reduce system aberrations **j)** from > 45 nm (top) to < 15 nm (bottom) root mean square error (RMSE), by iteratively measuring wavefronts derived from images of 500 nm beads and applying equal and opposite wavefronts via the deformable mirror. Left: wavefront representation; right: amplitudes for each Zernike mode (3-20). Representative axial views of 500 nm beads before **k)** and after **l)** correcting system aberrations are also shown. Images prior to correction show obvious asymmetries (red arrow, **k)**), which are largely removed post correction, resulting in more symmetric images with smaller axial (Z) extent **m)**, as quantified by full width at half maximum (FWHM). Scale bar in **k)**: 1 μm. Means and standard deviations in **m)** are derived from N = 25 beads.

We began by obtaining a benchmark calibration using a gold standard, commercially available Shack-Hartmann wavefront sensor (SHWFS, **Fig. 3b**). We applied positive and negative voltages to each DM actuator (**Fig. 3a**), imaging the associated fluorescence from a single 500 nm bead (the ‘guide star’) at each voltage onto the SHWFS. We then converted the measured SHWFS images into wavefronts by using the manufacturer’s software, then determined the linear relationship between wavefront and voltage, yielding the influence functions for each actuator (**Fig. 3b**). We empirically determined the illumination power and exposure time (1 s) necessary to produce high quality wavefronts, finding that with these settings we could reliably calibrate the mirror in 118 s using the SHWFS (115 s acquisition and 3 s for processing, **Fig. 3e**).

Our sensorless PD algorithm requires known diversity aberrations to sense an unknown wavefront. Given that our microscope also incorporated a piezoelectric stage, we reasoned that stage defocus could readily provide such known diversities and is an especially convenient choice as it does not rely on a calibrated mirror. For each actuator, we applied the same voltages as for the SHWFS calibration, but also applied different stage defocus values (± 2 μm, ± 1 μm, 0 μm) at each voltage and captured the associated images using our electron multiplying CCD (EM-CCD) detector (**Fig. 3c**). Feeding each set of five images into our PD algorithm produced a wavefront estimate for each voltage (**Methods**), subsequently enabling us to determine the induced wavefront as a function of voltage. Unlike the SHWFS, PD cannot directly estimate tip/tilt components in the wavefront. We thus developed an indirect method for estimating the contribution of these Zernike modes, relying on PD’s ability to estimate the object (and thus, the tip/tilt-induced displacement of the object at each voltage, **Methods, Supplementary Fig. 5**).

The tip/tilt corrected influence functions closely resembled the SHWFS ground truth (**Fig. 3d**). Even though the PD-derived influence functions required ten measurements (five defocus values x two voltages) per actuator instead of only two, we found that the greater sensitivity of our EM-CCD allowed us to acquire each image in only 30 ms, and with ~3-fold less power than when using the SHWFS. Thus, including both the shorter acquisition time (53 s) and computation time (11 s), the total time for calibrating all actuators was only 64 s, faster than the calibration with the SHWFS (**Fig. 3e**). We note that our PD-derived calibration time is also several hundred times faster than previous indirect methods^28,29^.

To compare the PD-derived calibration more quantitatively to that derived from the SHWFS, we conducted more detailed assays^27^ in which we inverted the influence functions to obtain control matrices, applied known Zernike aberrations (Zernike modes 1-20, each with amplitude 150 nm) and measured the degree to which the correct aberration was applied with the SHWFS (**Fig. 3f**). The SHWFS-calibration (**Fig. 3g**) provided near ideal performance (**Fig. 3i, Supplementary Fig. 6**, error 0 ± 7 nm, measured across on-target as well as off-target entries in our assay, N = 3543 measurements). Applying the same assay using the PD calibration produced a similar result (**Fig. 3h**), albeit with more noise (**Fig. 3i, Supplementary Fig. 6**, error -1 ± 12 nm, N = 3581). Given that the PD calibration used ~20-fold less fluorescence than the SHWFS, it is perhaps unsurprising that it produced a noisier result. We found that using alternate diversity phases (e.g., astigmatism, **Supplementary Fig. 7**) improved in the PD response, suggesting that the choice of diversity may also influence PD performance in this assay. Incorporating tip/tilt modes into our calibration was essential, as characterization assays without tip/tilt compensation were considerably worse (**Supplementary Fig. 5**). We also found that PD yielded calibrations of similar quality on an extended sample consisting of multiple 500 nm beads (**Supplementary Fig. 8**), which was impossible with the SHWFS calibration scheme as implemented here, since it required a single bead.

Perhaps most importantly, we investigated whether the PD-derived calibration could remove system aberrations by ‘flattening’ the wavefront measured from our beads, a key prerequisite for successful AO. To this end, we first used PD to estimate the wavefront from single 500 nm beads (using the same defocus diversities as for the calibration), finding small but non-negligible aberrations (**Fig. 3j**, typical RMS wavefront error of 46 nm measured over the first 20 Zernike modes). These aberrations also manifested as slight asymmetries in bead images, evident in axial views (**Fig. 3k**). Next, we implemented a feedback scheme whereby we used our PD calibration to induce an equal and opposite wavefront, thereby reducing the aberration (**Methods**). This method reliably reduced the wavefront error to less than 15 nm (**Fig. 3j**) within a few iterations, resulting in more symmetric bead images (**Fig. 3l**) and submicron axial extent (**Fig. 3m**).

### Aberration correction on extended samples using phase diversity

Having demonstrated that phase diversity may be used to calibrate a deformable mirror, we next turned to aberration correction on extended samples consisting of multiple 500 nm fluorescent beads (**Fig. 4**). We introduced test aberrations *ϕ*_Abe_ of varying composition and magnitude, collected additional images with added diversity aberrations, and evaluated the extent of our algorithm to sense and correct the test aberrations after one or more cycles of wavefront sensing and correction (**Fig. 4a, Methods**).

**Fig. 4.**
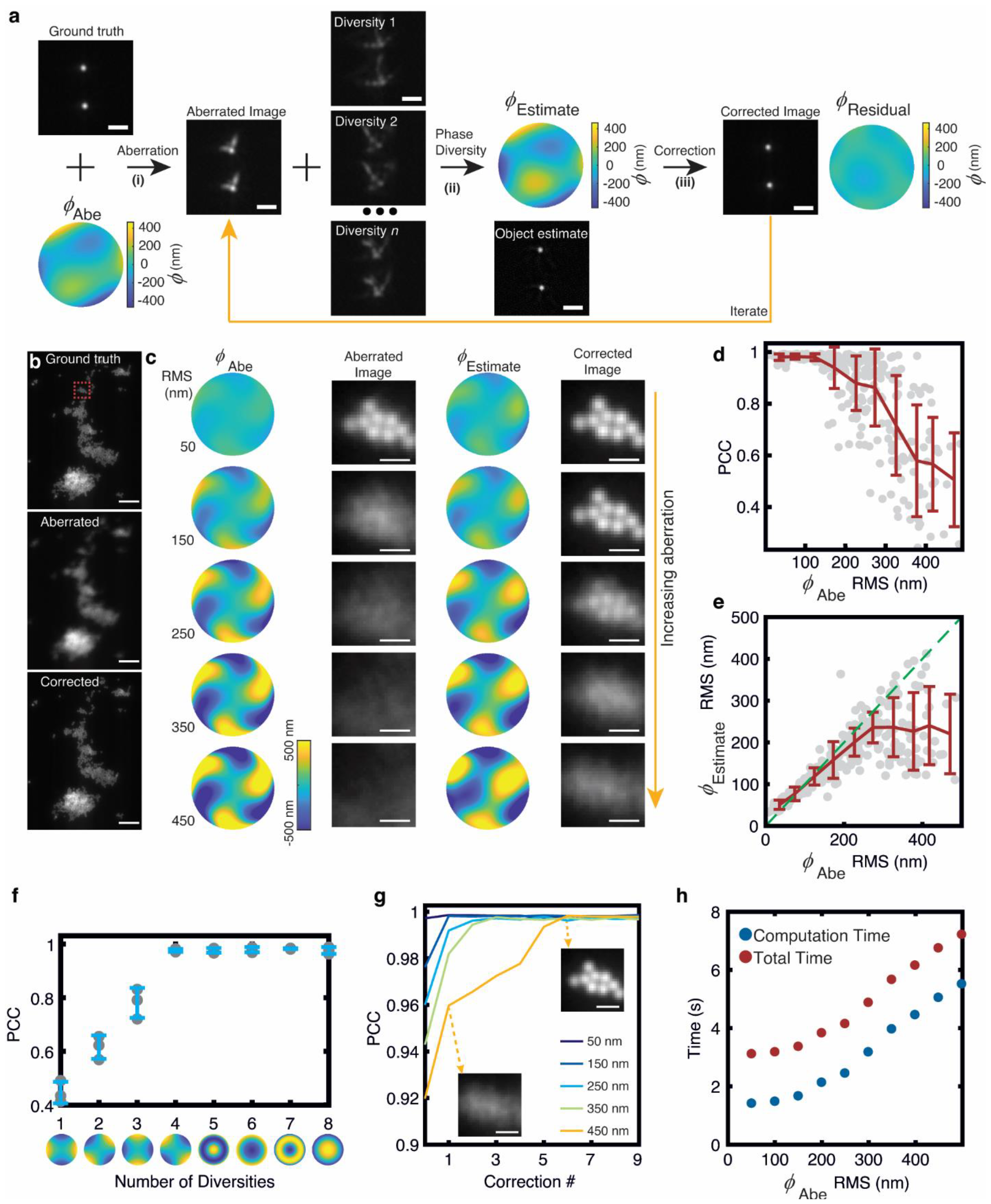
Phase diversity-based AO on extended, multi-bead samples. **a)** We introduce known aberrations (*ϕ*_Abe_) to a multi-bead sample and attempt to sense and correct these aberrations using phase diversity. We (i) collect a series of diversity images (Diversity 1, Diversity 2,… Diversity *n*) which are then used to (ii) estimate the unknown wavefront (*ϕ*_Estimate_) and object using phase diversity. The deformable mirror is then programmed to (iii) apply an equal but oppositely signed wavefront, providing a corrected image. We then re-measure the wavefront (*ϕ*_Residual_) and iterate if necessary. **b)** Example images of 500 nm beads with system aberrations corrected (ground truth, top), after inducing aberration with RMS wavefront distortion 150 nm (middle), and after one cycle of phase diversity-based AO correction using four diversity images (bottom). **c)** Effect of increasing aberrations. Columns from left: same test aberration as used in **b)** with indicated RMS wavefront distortion; higher magnification view of associated aberrated subimage, corresponding to red dashed rectangle in **b)**; estimated wavefront produced by phase diversity; corresponding image after correction is applied. Rows show effect of increasing the RMS amplitude of *ϕ*_Abe_. These results were obtained with a single cycle of wavefront estimation and correction. **d)** Assessing correction using Pearson correlation coefficient (PCC, y axis) when applying random assortment of *ϕ*_Abe_ with indicated wavefront distortion (x-axis). Results are shown for a single cycle of wavefront sensing and correction; PCC is measured relative to ground truth images. Individual trials (N = 315) are shown as scatterplot, mean and standard deviations are also shown, binning points in 50 nm increments. Data are pooled from three different fields of view. **e)** As in **d)** but assessing wavefront estimation by examining RMS wavefront distortion in estimated aberration vs. test aberration. Dashed green line indicates values in which the input and estimated RMS values are identical. **f)** Quality of images resulting from single correction (as assessed by PCC, y-axis) as a function of the number of diversity phases (x-axis) used in wavefront estimation. Here the test aberration was the same shape as in **b, c)**, with 236 nm RMS distortion. Individual data points as well as means and standard deviations are shown from three independent imaging fields. Diversity phases (±1 µm amplitude oblique astigmatism, horizontal astigmatism, defocus, and spherical aberration) are shown below the x-axis; data shown in **b-e, g, h)** all use the first four phases. **g)** Quality of correction (PCC, y-axis) as a function of number of correction cycles (x-axis) for *ϕ*_Abe_ with 50 nm, 150 nm, 250 nm, 350 nm, 450 nm wavefront distortions using same data as in **b, c)**. Insets with dashed arrows show first and sixth correction with 450 nm RMS wavefront distortion in *ϕ*_Abe_. **h)** Total time (red) and computational time (blue) for first correction, as a function of RMS magnitude of *ϕ*_Abe_. Computational time scales nonlinearly as a function of RMS magnitude due to the increased number of iterations, whereas acquisition, file saving, and instrument overhead contribute a fixed time cost (here 1.7 s). Bead crops in **c, g** are scaled to the same intensity range. Scale bars: 2 μm **a, c**, insets in **g**), 5 μm in **b**).

On extended, densely labeled imaging fields, a single cycle using the aberrated image and four diversity images (±1 µm oblique astigmatism; ±1 µm vertical astigmatism) proved sufficient to restore images contaminated with up to ~200 nm RMS wavefront distortion (**Fig. 4b, c**). At this level of degradation, individual beads were severely blurred and could not be easily discerned in the input images, yet correction resulted in the same clarity as ground truth images without system or added aberrations. As expected, increasing the magnitude of aberrations caused progressive deterioration in the quality of our correction, yet we observed noticeable improvement in signal even when the aberrated images were so blurred that no hint of bead structure remained (**Fig. 4c**).

We found the Pearson correlation coefficient (PCC) between corrected and ground truth images useful in characterizing correction performance over a variety of test aberrations (**Fig. 4d**). PCC values above ~0.9 usually were associated with corrections that provided clear improvement over the aberrated input, and we found that most data collected with test aberrations with < 250 nm RMS wavefront distortion fell into this category. PCC values below 0.9 usually indicated obvious aberration. Although in some cases we obtained a PCC > 0.9 even for test aberrations with RMS wavefront distortion exceeding 300 nm, in general PCC values exhibited a noticeable ‘shoulder’ at ~250 nm RMS distortion, with an increasingly steep drop and spread in PCC extent below this shoulder.

To better understand this behavior, we also examined *ϕ*_Estimate_, the wavefront estimate produced by phase diversity (**Fig. 4c, e**). If the test aberration *ϕ*_Abe_ was being accurately sensed, we would expect *ϕ*_Estimate_ to closely resemble *ϕ*_Abe_. When examining the RMS distortion in both quantities, we found that they agreed well until ~250 nm, where we again observed a ‘shoulder’ beyond which we observed an increasing difference between the RMS values of *ϕ*_Estimate_ and *ϕ*_Abe_ (**Fig. 4e**). These results suggest that inaccurate wavefront sensing is a primary reason that correction fails past ~250 nm RMS test aberration.

Next, we investigated methods to improve performance past this limit. First, we explored varying the magnitude of the diversity phases. In general, we found that larger diversity magnitudes offered better sensing, although in all cases sensing still degraded in the presence of an increasing magnitude of test aberration (**Supplementary Fig. 9**). Second, we varied the number and type of diversity phases, using different Zernike modes as the different diversities (**Fig. 4f**). Using a test aberration with 250 nm RMS wavefront distortion, we found that performance improved up to four diversity phases (the number used thus far, **Fig. 4b-e**) and then plateaued, at least for the choice of diversities we tried here. Further testing of different diversities (e.g., random phases) might offer improved performance with fewer diversities. Interestingly, we also confirmed the prior result from simulations (**Supplementary Fig. 3**), whereby coma and trefoil Zernike modes proved ineffective for wavefront sensing. Finally, we explored the effect of multiple cycles of wavefront sensing and correction (**Fig. 4g**). This strategy proved the most effective, as we were able to fully correct larger aberrations (up to 450 nm RMS distortion, the largest value we tested) simply by increasing the number of correction cycles.

The time required for a single cycle of wavefront sensing and correction depended on the magnitude of test aberration, with larger aberrations requiring more time (**Fig. 4h**). For test aberrations with RMS wavefront distortions < 250 nm, the computational burden associated with wavefront sensing was on par with acquisition related timing (1.7 s, including file saving and hardware-related delays, see **Methods**). For larger aberrations, the computational burden was several-fold larger. Nevertheless, given anticipated improvements in both data acquisition and algorithmic efficiency, we are confident that the second-level time resolution we report for our imaging field (512 × 512 pixels, ~53 × 53 μm^2^) could be substantially reduced in the future.

### Phase diversity-based adaptive optics restores cellular images contaminated with aberrations

Finally, we investigated the use of phase diversity-based wavefront sensing for AO correction on cellular samples (**Fig. 5**). On fixed U2OS cells immunostained for microtubules (**Fig. 5a, e**), aberrations with 219 nm RMS wavefront distortion blurred images to the extent that individual fibers were indiscernible (**Fig. 5b, f**). We used PD to sense the aberrated wavefront, finding that we could dramatically improve image quality with only two correction cycles, largely restoring the appearance of microtubule fibers (**Fig. 5c, g**). However, close inspection of the corrected result also revealed that correction was incomplete (compare **Fig. 5e, g**). We suspect the reason for this is mainly due to the out of focus fluorescence resulting from widefield illumination and detection on these 3D samples. As our PD algorithm does not currently account for the 3D nature of the object, it has no way to model such background, and likely misinterprets it as out of focus background from the focused image plane as well as other aberrations. In support of this hypothesis, we were able to partially compensate for such effects by intentionally neglecting to correct the defocus Zernike mode (**Fig. 5d, Methods**). The resulting AO correction without defocus produced better restoration than the full AO correction (**Fig. 5h**), albeit still worse than the ground truth result (**Fig. 5e**).

**Fig. 5.**
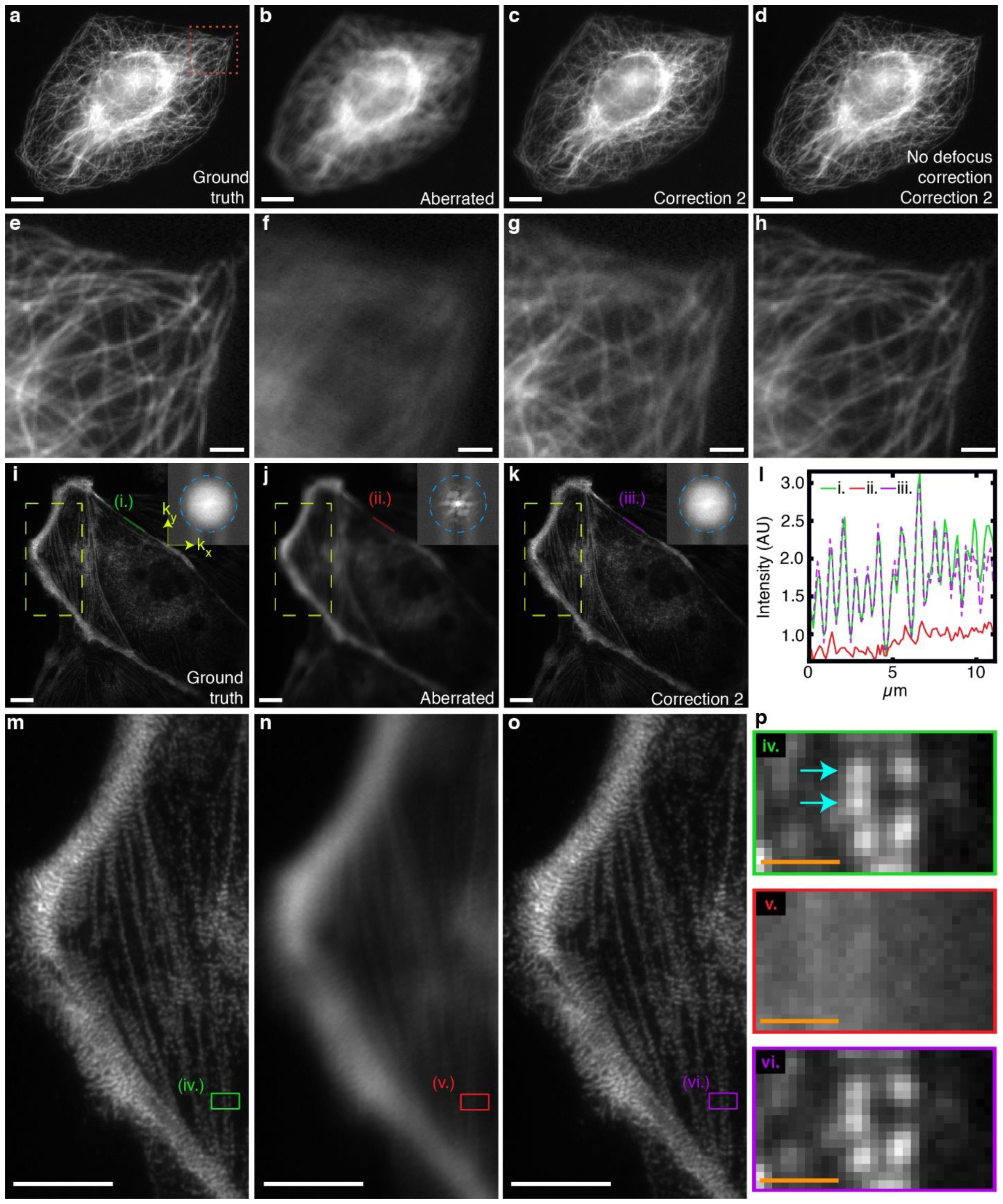
Phase diversity-based AO on biological samples. **a-d)** U2OS cells were fixed and immunostained for microtubules. Images of ground truth **a)**, after aberration with 219 nm RMS wavefront distortion **b)**, after resulting AO correction (second correction cycle) **c)**, and AO-corrected but without defocus correction (also second correction cycle) **d)** are shown. **e-h)** Higher magnification views corresponding to the red dashed rectangular region in **a). i-k)** Ptk2 cells were fixed and immunolabeled against myosin heavy chain. Images of ground truth **i)**, aberrated with 311 nm RMS wavefront distortion **j)** and corrected after measuring wavefront distortion with phase diversity **k)** (result corresponds to second correction cycle, see also **Supplementary Fig. 10**). Insets show Fourier transform magnitudes (displayed after logarithmic transformation), with dashed circle indicating 1/300 nm^-1^ spatial frequency. **l)** Line profile comparison along green (i), red (ii), and magenta (iii) lines in **i-k)**, showing that AO correction recovers peak profiles which are absent in the aberrated image. **m-o)** Higher magnification views of dashed rectangular regions in **i-k). p)** Higher magnification views of green (iv), red (v), and magenta (vi) rectangles in **m-o**). Cyan arrows highlight puncta separated by ~300 nm that are resolved in ground truth images, absent in aberrated image, and recovered in AO corrected image. Scale bars: 10 μm **a-d, i-k), m-o);** 2 μm **e-h)**; 1 μm **p)**. Images have been flat-fielded (**Methods**).

We were nevertheless encouraged by the performance of our current 2D PD algorithm on these cellular volumes, as it suggests that PD will likely perform even better on microscope systems with optical sectioning. As another test case more suited to the widefield microscope employed here, we examined immunostained myosin in fixed Ptk2 cells (**Fig. 5i**). As these samples were much thinner (generally of submicron thickness), they provided a closer approximation to a 2D biological sample than the U2OS cells. Myosin is also known to assemble into bipolar filaments of length ~300 nm in vitro^30^ and in cells^31^, providing an additional test of the resolution of our system.

In ground truth images, we resolved periodic striations of myosin along stress fibers (**Fig. 5i, l, m**), with individual puncta sized ~300 nm or above. While we could not universally resolve the separation between bipolar myosin heads, Fourier transforms revealed spectral density out to ~300 nm (**Fig. 5i**), and we occasionally found puncta separated by ~300 nm (**Fig. 5p**). By contrast, introducing aberrations with 311 nm RMS wavefront distortion (**Fig. 5j**) reduced spatial resolution to the point that myosin striations (**Fig. 5l, n**) or individual puncta (**Fig. 5p**) were completely absent. A single cycle of PD-based AO correction largely restored imaging performance (**Supplementary Fig. 10**), and after two cycles we could no longer discern differences between the ground truth and AO-corrected images (**Fig. 5k, l, o, p**).

## Discussion

In samples thicker than a single cell, the presence of optical aberrations hinders diffraction-limited imaging. AO provides a means of counteracting such aberrations, yet despite ongoing refinement, these methods remain the province of relatively few labs. We believe broad dissemination of AO has been hampered for several key reasons. First, additional hardware must be integrated into the optical path, which is not trivial to set up, and often adds considerable complexity and cost relative to the base microscope. In this regard, phase diversity (along with many other indirect methods) requires the addition of an adaptive element and accompanying relay optics only, simplifying microscope design, and improving emission-side light efficiency compared to direct sensing approaches. Second, AO methods require dedicated algorithms and software for wavefront sensing and corrective feedback, which until recently have not been freely available^27^. Finally, many AO methods (especially indirect) require substantially more measurements (or time) to sense the aberrations than to acquire the aberrated image. We think phase diversity is highly promising in this regard, as only a few additional images are required to sense the wavefront, and sensing can occur on sub-second timescales.

As we demonstrate when calibrating a deformable mirror (**Fig. 3**), our existing implementation of phase diversity is orders of magnitude faster than related indirect approaches^28,29^, and also provides faster calibration than direct sensing using a commercially available SHWFS. We also show that our method can correct severe aberrations induced in biological samples (**Fig. 5**), recovering diffraction-limited performance. We believe these results pave the way for a future generation of adaptive microscopes that do not rely on SHWFS, minimize instrument complexity, and provide AO capability rapidly and with minimal extraneous dose or loss in temporal resolution.

Our method can be improved, and its scope extended beyond the proof-of-concept studies demonstrated here. First, the 3D object is not modeled in our current 2D phase diversity method. Three-dimensional phase diversity^24^ provides a more realistic imaging model and might improve the accuracy of wavefront sensing and subsequent correction on 3D biological samples, particularly if there is substantial out-of-focus light (**Fig. 5a-h**). Second, more accurately modeling the PSF^32^ or noise (e.g., Poisson^23^ instead of Gaussian, as assumed here) might similarly improve both wavefront sensing and object estimates. Third, here we considered only a relatively small field of view, over which aberrations were isoplanatic. Imaging larger, more heterogeneous samples will likely require tiling, with sensing and correction (or at least differing object and phase estimates) applied for each tile^33,34^. Fourth, the question of which diversity aberrations are best applied to optimally sense an aberrated wavefront is an open one^35^, and deserves further study. Fifth, although we have focused here on deformable mirror calibration and aberration correction, our phase diversity method is likely to facilitate other methods that rely on wavefront sensing, such as remote refocusing^36,37^. Sixth, approaches that use neural networks^38^ in conjunction with additional diversity images^25,39^ for wavefront sensing may offer improved performance in highly aberrating tissue^40^, but are still quite slow compared to classical methods like ours. An intriguing direction might be to combine these approaches, offering improved speed and performance relative to that offered by either class alone. Seventh, the speed of our algorithm could likely be further improved if the current MATLAB implementation was rewritten in a compiled programming language, and perhaps if a different optimization method were used. Finally, a clear next direction is the application of our method to systems that provide optical sectioning, including light sheet and confocal microscopy.

## Methods

### Sample preparation

To adhere 500 nm diameter yellow-green fluorescent beads (ThermoFisher Scientific, F8813) to #1.5 coverslips (Thorlabs, CG15XH), we (i) cleaned coverslips with methanol; (ii) pipetted ~5 μL Poly-L-Lysine solution (Millipore Sigma, P4832-50ML) onto the center of the cleaned coverslips; (iii) waited ~5 minutes and rinsed the coverslips with methanol; (iv) allowed coverslips to dry; (v) diluted 500 nm beads ~10,000-50,000-fold in methanol; (vi) pipetted diluted bead solution onto the center of the coverslips. Beaded coverslips were then stored dry until use.

Ptk2 cells were grown in Eagle’s Minimum Essential medium (ATCC, 30-2003) supplemented with 10% Fetal bovine serum (Gibco, 26140079), in a 5% CO_2_ incubator. They were dissociated with 0.05% Trypsin-EDTA (Gibco, 25300054) and replated on #1.5 coverslips (Warner Instruments, 64-0734). Cells were kept in the incubator for 24 hr, before being fixed in pre-warmed (5 min in 37° C water bath) 4% formaldehyde (Electron Microscopy Sciences, 15710) in PBS solution for 5 min and permeabilized in pre-warmed (to 37° C) 0.2% Triton X-100 (Sigma, T8787) in PBS for 5 minutes. After a quick rinse with PBS, cells were incubated in 1mg/mL L-Lysine (Sigma, L5501) + 1% IgG free BSA (Biotium, 22013) in PBS solution for 15 min at room temperature. For staining, cells were incubated with mouse anti-myosin heavy chain Clone H11, IgM (Biolegend, 922704) and Rabbit anti Phospho-Myosin Light Chain 2 (Thr18/Ser19) (Cell Signaling Technology, 3674), both at 1:200 dilution in PBS containing 1% BSA. Samples were incubated in primary antibodies at 37° C for 1 hour, washed once (with 5 min incubation) in PBS containing 0.05% Tween-20 (Fisher BioReagents, BP337-500), and washed twice in PBS (5 min incubation each time). Secondary antibody staining was performed at 37 ° C for 1 hour, using a solution containing 1% IgG free BSA in PBS, goat anti-rabbit IgG secondary antibody labeled with Alexa Fluor 568 (ThermoFisher Scientific A-11011) at 1:50 dilution, and goat anti-mouse IgM secondary antibody labeled with Alexa Fluor 488 (ThermoFisher Scientific A-21042) at 1:400 dilution. Cells were incubated in PBS containing 0.05% Tween-20 for five minutes, then incubated twice for five minutes in PBS. Finally, immunostained cells were mounted in ProLong Gold, Antifade Mountant (Thermo Fisher, P10144), cured in the dark at room temperature for 24 hours, and stored at -20° C until imaging.

U2OS cells were fixed and immunostained for microtubules as previously described^41^.

### Widefield adaptive optics system

To demonstrate our phase diversity wavefront sensing method and compare its performance to Shack-Hartmann based wavefront sensing, we constructed a widefield system with deformable mirror and Shack-Hartmann sensor in the emission path.

All optics and optomechanics were bolted to a 3’x5’ table (TMC,784-636-02DR and 12-416-84) floated with room air. To excite fluorescence, the beam from a 488 nm laser (Coherent, Sapphire 200 mW, 1137967) was expanded 15x (Thorlabs, GBE15-A), reflected using a dichromatic mirror (DC, Semrock, Di03-R488-t3-25×36), and focused with lens L1 (Thorlabs, 250 mm focal length achromat, AC254-250-A-ML) at the back focal plane of a 60x, 1.2 NA water immersion objective (OBJ, Evident Scientific, 1-U2B893) held in a rapid automated modular microscope (RAMM) system equipped with motorized (for coarse axial objective positioning and lateral sample positioning) and piezoelectric (150 micron travel, for fine axial sample displacement) stages (Applied Scientific Instrumentation, RAMM-BASIC and PZ-2150-FT). We used an automated beam shutter (Thorlabs, SH05R) and neutral density filters (Thorlabs, NE20A-A) for beam shuttering and attenuation as needed.

Fluorescence was collected in epi-mode through the same objective and was transmitted through L1 and DC. A 500 mm focal length achromat (L2, Thorlabs, ACT508-500-A-ML) placed in 4f relation to L1 served to magnify and image the objective’s pupil 2x (from 7.2 mm to 14.4 mm), thereby serving to nearly fill the ~15 mm diameter of our deformable mirror (DM, Imagine Optic, MIRAO 52ES), placed at ~5 degree incidence to minimize off-axis aberrations. Post DM, a 750 mm focal length achromat (L3, Thorlabs, ACT508-750-A-ML) placed one focal length from the DM served to image the sample plane onto an electron multiplying charge coupled device (EM-CCD, Oxford Instruments, iXon Ultra 888, DU-888U3-CS0-#BV). The pixel size in image space, S_px_, was 104 nm (= 13 μm/[(250/3)*(750/500)]) and the illuminated area spanned the majority of the ~106.5 μm field of view. We placed a bandpass filter (EF, Semrock, FF03-525/50) upstream of the camera to reject pump light and water-cooled the EM-CCD to minimize fan induced vibrations (Solid State Cooling Systems, ThermoCube, 10-400-1C-1-AR).

For some experiments, we diverted the fluorescence to a Shack-Hartmann wavefront sensor (SHWFS, Imagine Optic, HASO4 First Shack-Hartmann Wavefront Sensor, IO-6WFS201) with a flip mirror (FM, Newport, 8893-K) placed post-L3, prior to the EM-CCD. A 180 mm focal length achromat (L4, Thorlabs, AC508-180-A-ML) placed in 4f relation to L3 served to demagnify and relay the image of the objective pupil from DM to the Shack-Hartmann sensor. When using the Shack-Hartmann wavefront sensor, we also placed an iris at the intermediate image plane (the focal point midway between the L2/L1 lens pair) to isolate the signal from a single 500 nm diameter fluorescent bead, and placed a bandpass emission filter (EF, Semrock, FF03-525/50) in front of the sensor. These optics are shown in **Supplementary Fig. 4**.

Bead studies employing phase diversity used a laser power of 180 μW, measured by placing a power sensor (PM100D Console, S170C Sensor, Thorlabs) after the objective. Bead studies employing the SHWFS used 524 μW, and cell imaging studies 1.07 mW.

### Microscope control software and hardware

The microscope was controlled using a custom-designed program in LabVIEW Version 2023 Q1 (National Instruments) which calls the Andor SDK V2.104.30084.0 (for control of the EM-CCD camera) and the Imagine-Optic WaveKit V4.3.2 software (for control of the Shack-Hartmann sensor and deformable mirror). The LabVIEW program also controls a Piezo stage (PZ-2150-FT, ASI), laser shutter (SH05R, Thorlabs), and motorized flipper mount (8893-K, Newport). The control software was run on a Colfax ProEdge™ TRX4400 Windows 10-based workstation PC featuring an AMD Ryzen Threadripper PRO 3995WX 64-core CPU, 256 GB system RAM and an Nvidia RTX A5000 with 24 GB video RAM. This computer was also used for all computational processing.

### Deformable mirror voltage offset

Significant aberration is introduced when the deformable mirror has no voltage applied. A voltage offset can be applied to cancel these and other sources of system aberrations, a process we refer to as “flattening” the mirror. This offset then serves as the base set of voltages to which all other voltages are added to. Such offsets were used for all experiments. An initial set of voltages is typically provided by the DM manufacturer, but this offset does not account for any other sources of system aberration or longer-term drift. An updated flat was generated using the AO correction loop procedure described below.

### Phase Diversity algorithm

The goal of the phase diversity algorithm is to take the input diversity images, their corresponding known diversity wavefronts, and parameters of the system (including numerical aperture, 1.2; image pixel size, 104 nm; and central emission wavelength 532 nm) and return an estimate of the unknown wavefront. We implement this algorithm in MATLAB.

When performing on-line wavefront estimation, LabVIEW initiates the algorithm using a MATLAB script node which calls the MATLAB function `processPhaseDiversityImages`, a wrapper function which passes the inputs from LabVIEW to MATLAB and calls the function chain to implement the phase diversity algorithm. This set of functions performs a series of pre-processing steps, passes the resulting inputs to the algorithm, and returns the wavefront and object estimates as outputs which are passed back to LabVIEW.

First in this chain is the function `reconstructZernikeAberrations`, which initiates data pre-processing and passes this result as input to the algorithm function. Image pre-processing is handled by `fileIO_lvtiff2mat`, which transfers the saved EM-CCD camera images from disk into MATLAB. The full-frame images are loaded from disk, cropped to the intended size for processing, a constant background value is subtracted, and an edge-tapering applied using a Gaussian smoothing kernel with size 10×10 pixels and standard deviation of 3 pixels.

Additional pre-processing steps are performed by the function `zernretrieve_pre`. First, the images passed from `fileIO_lvtiff2mat` are Fourier transformed. Second, the Zernike polynomial functions for the modes to be estimated are computed. Finally, wavefronts corresponding to the diversity phases are constructed from their known coefficients. The pre-processed data and parameters are then passed as inputs to the algorithm function, `zernretrieve_loop`.

The algorithm uses Gauss-Newton optimization to find the wavefront that minimizes an objective function *J* (**Supplementary Note 1**). In frequency space, it is given in terms of the optical transfer functions (OTFs, *S*_*k*_), modulation transfer functions (MTFs, |*S*_*k*_|), and frequency-space diversity images (*D*_*k*_):

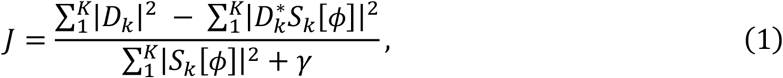

with *γ* being a regularization term set to 1×10^−6^ (determined from simulations, **Supplementary Fig. 2b**) and * denotes the complex conjugate.

We express the wavefront in a basis of *M* Zernike polynomials and update the Zernike coefficients **c** by computing the gradient, **g**, (an M x 1 vector) and the pseudo-Hessian, **H**, (an M x M matrix) of the Gaussian likelihood function. The update equation from iteration *q* to iteration *q* + 1 is given by:

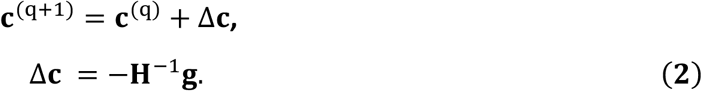

In practice, the wavefront is initialized by setting all coefficients to zero **c**^(0)^ = 0.

We evaluate J for convergence by comparing the marginal change in *J* to the total change since iteration began. Further iteration is terminated if this quantity is less than 0.001, if J begins increasing, or if the maximum number of iterations (100 in all tests) is reached. The change in J is given by

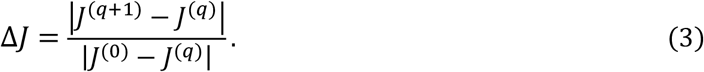

To construct the gradient, we evaluate

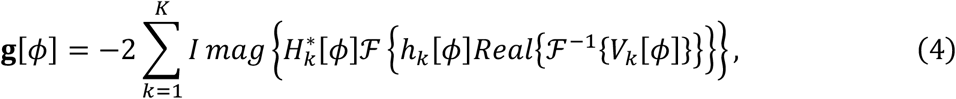

where

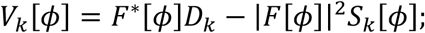

ℱ and ℱ ^−1^ denote the Fourier transform and its inverse, respectively; *H*_*k*_(*ϕ*) and *h*_*k*_[*ϕ*] denote the pupil function and its inverse Fourier transform, respectively; and *F*[*ϕ*] is the Fourier transform of the object.

To reduce the dimensionality and obtain the gradient in terms of a reduced set of Zernike coefficients, we take the inner product between this gradient and the Zernike polynomials to produce an Mx1 gradient vector.

The Hessian **H** is constructed by calculating the elements of the matrix as

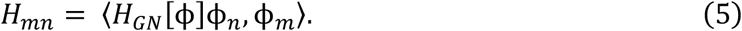

Following ref.^22^, this expression can be evaluated as

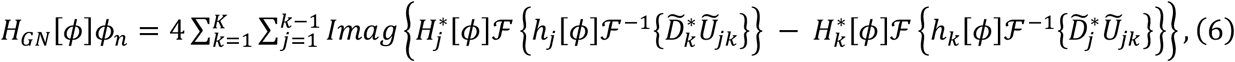

where

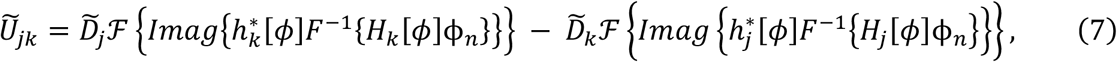

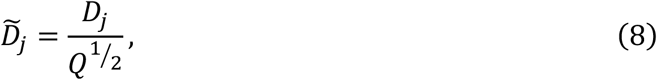

and

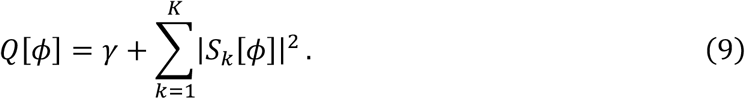

To speed up this computation, we performed the calculations on our GPU using MATLAB’s `gpuArray` functionality. Further, we optimized the calculations by avoiding looping over the sums, but instead calculating the terms in Eq. 6 for all indices in parallel. As the outer sum is taken from *k* = 1 to *K*, and the inner sum from *j* = 1 to *k* − 1, we first determined all required index pairs and then calculated the term for the required pairs only.

To further increase the computation speed, we implemented the calculation of the Zernike polynomials as a handle class, i.e., all required Zernike polynomials are calculated only once and then cached, avoiding duplicated computation. The aberrations in each iteration are then computed by multiplying the new coefficient estimates with the precomputed Zernike polynomials.

Finally, the coefficients are updated using the previous coefficients, the Hessian, and the gradient using Eq. 2. These steps are shown in schematic form in **Supplementary Fig. 1**.

Our implementation can estimate a wavefront in ~100 ms (e.g., for the 128 × 128 crops around a single bead used for DM calibration, **Fig. 3**), with time cost generally increasing in proportion to the image area (e.g., on the seconds level for the first correction cycle in the multi-bead tests shown in **Fig. 4h**). We also found that computational cost increased as the magnitude of wavefront distortion increased, which was due to the increased number of required iterations.

### Phase diversity-based calibration of the deformable mirror

The deformable mirror calibration procedure was implemented using the custom LabVIEW program described above. To acquire the control matrix, negative and positive voltages (±0.03 V) were applied to a single actuator and image stacks were acquired with axial stage positions ±2 μm, ± 1 μm, 0 μm, with each axial defocus used as a diversity phase. Images were exposed for 30 ms for each voltage value at each stage position, and overall acquisition time was minimized by moving the stage during the camera readout period, resulting in a total time of 1.03 s per actuator, including moving the stage, exposing, readout, and switching mirror voltages. This process was repeated for each of the 52 mirror actuators. After acquisition, the influence functions for all 52 actuators were computed off-line using MATLAB.

To reduce readout time, images were acquired with a cropped sensor mode of 512×512 pixels. For DM calibrations featuring a single bead (**Fig. 3**), images were further cropped to a size of 128×128 pixels to minimize processing time. For calibrations featuring multiple beads the full 512×512 acquisition size was used.

Off-line phase diversity processing was performed using the script `script_Calibration`. When using stage defocus, we computed the high NA defocus phase^32,42^ corresponding to each stage position *Z*:

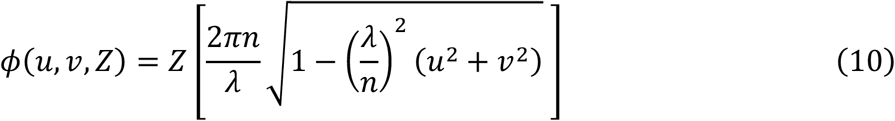

where *u,v* ranges from 0 to 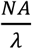, *n* is the refractive index and *λ* is the emission wavelength; and used these phases as the diversity phases in `reconstructZernikeAberrations,` subsequently estimating the wavefront as described above. After estimating the wavefront for each voltage for a given actuator, the influence function for each of the 52 actuators were computed from the slopes of each coefficient value given by

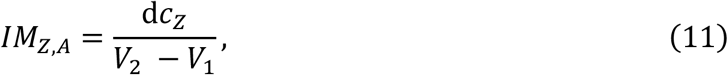

where *IM*_*Z,A*_ is the interaction matrix slope value representing the change in Zernike coefficient d*c*_*Z*_for mode Z, as DM actuator A changes voltage from V_1_ to V_2_. In all cases here we used V_1_ = - 0.03V and V_2_ = +0.03V. After all values were computed the control matrix was obtained by pseudo-inverting the interaction matrix using the MATLAB function `pinv` with a tolerance value of 0.005.

### DM calibration with alternative diversity phases

In some experiments, we calibrated the DM using astigmatism as a diversity phase (**Supplementary Fig. 7**), finding that this choice gave lower off-target noise than our stage defocus-based calibration. In this case the applied astigmatism commands were derived from a previous calibration, and we used four permutations of horizontal or oblique astigmatism with values of ± 1 μm as our diversities. In addition to the offset voltage and diversity phase voltages, we added additional voltages (−0.03V/+0.03V) to the actuator being measured. Wavefronts were measured using the on-line wavefront estimation procedure described above. Off-line processing was performed to obtain the influence functions from the slope of the individual wavefronts for each actuator and to calculate tip/tilt from the object estimates as was done in the standard PD DM calibration procedure.

### Estimating tip and tilt

Tip and tilt are first-order Zernike modes which represent vertical and horizontal translations in the image plane. These wavefront components are not accounted for by the phase diversity algorithm but are critical to include in DM calibration to ensure that image translations are not inadvertently included when applying commands. To estimate residual tip/tilt that arise when different voltages are applied to the DM, we rely on object estimates produced by PD, whereby relative differences in tip/tilt manifest as displacements in object position resulting from the different voltages.

To estimate displacement between object estimates, we compute their cross-correlation with sub-pixel accuracy using a single-step discrete Fourier transform algorithm (ref.^43^, https://www.mathworks.com/matlabcentral/fileexchange/18401-efficient-subpixel-image-registration-by-cross-correlation, **Supplementary Fig. 5a**). The computed shift in each axis *pj* is given in units of pixels, which must be converted to units in pupil space to be incorporated into our control matrix. To determine the appropriate conversion factor, we simulated object estimates with different magnitudes of tip/tilt. Computing the translation as a function of tip/tilt Zernike coefficient amplitude yields a conversion factor *δ* of 0.8352 μm image space/µm coefficient amplitude. *δ* can then be used to convert *p*_*j*_ to coefficient amplitudes via

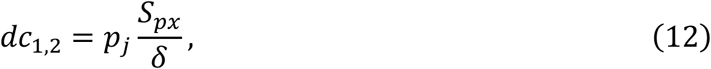

where *dc*_1,2_ are the relative Zernike coefficient amplitudes for tip/tilt (units of μm) and *S*_*px*_ is the size of one pixel (units μm).

This computation is performed when calibrating the DM, and when using phase diversity to validate a calibration using the characterization assay (**Supplementary Fig. 5d, e**). For the DM calibration, coefficient amplitudes as a function of actuator voltage are obtained for each of the 52 actuators using the object estimates obtained for each DM voltage pair. The slope value (units μm coefficient amplitude/V) that is used in the interaction matrix, *IM*_(1,2),*A*_ is given by:

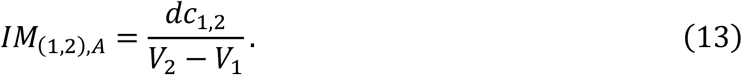

When validating the calibration using phase diversity (**Supplementary Fig. 5e**), we find the relative displacements between an object estimate arising from the issued command and an object estimate associated with a reference wavefront without the command. Eq. 13 is then applied to obtain the relative tip/tilt.

### Shack-Hartmann operation

The SHWFS was controlled via the custom LabVIEW program described above using WaveKit v4.3.2 SDK. For all experiments, the device was operated with an exposure time of 1 s. The processing pipeline to go from image data to Zernike coefficients was performed on-line using the Imagine Optic Wavekit software. First, the image data of the imaged bead was converted to wavefront slopes. Second, using the calibration data supplied with the device the slopes are converted to Zernike coefficients using the modal Zernike reconstruction parameter which outputs Zernike coefficients under the Wyant convention. To facilitate conversion to ANSI notation, we output 32 coefficients. Notation conversion was performed using custom LabVIEW code and the resulting ANSI coefficients were cropped to the desired set of 20, including tip and tilt.

The procedures for performing a DM calibration and its validation were identical to the phase diversity version, except that instead of acquiring defocus steps and computing wavefronts using the PD algorithm, wavefronts were acquired directly using the SHWFS as described above.

### Control Matrix Validation

We used a characterization assay to quantitatively assess calibration performance for Zernike modes 1-20. Voltages corresponding to a 0.15 µm amplitude of each tested mode were added to the offset voltage and applied to the DM. Defocus diversity image stacks were acquired using the same axial defocus positions (±2 μm, ±1 μm, 0 μm), exposure time (30 ms) and image crop size (128 × 128) as for the DM calibration. Wavefront estimates from the resulting diversity defocus image stacks were obtained as for the calibration data. The phase diversity algorithm estimates for modes 3-20 and tip/tilt were computed from the object estimate as described above. As the wavefront measurement may contain nonzero values corresponding to residual aberration, we acquired an additional reference wavefront by applying only the offset voltage to the DM and measuring this residual aberration. This reference wavefront was subtracted from each of the tested mode wavefronts to obtain the wavefront corresponding to the applied command for each mode.

When assessing calibration performance using the SHWFS, we applied modes 1-20 as above, then measured the resulting wavefront using the SHWFS. A reference wavefront with only the offset voltage applied to DM was subtracted from the result using the Imagine Optic SDK.

For each applied mode, the error was determined based on the target value for the Zernike coefficients. For example, to validate mode 3 (oblique astigmatism), the expected Zernike coefficient amplitude was 0.15 μm for oblique astigmatism and 0 for all other coefficients. We computed the error for every element in the matrices in **Fig. 3g, h** by subtracting the target coefficients from the measured coefficients.

The control matrix generated by the calibration is sensitive to the orientation of the measurement device. As our EM-CCD camera and SHWFS were mounted orthogonally with respect to each other, this change in orientation must be accounted for when validating calibrations which were originally acquired with a different modality. To perform a SHWFS validation of a control matrix generated using PD we first rotate the measured PD wavefronts by **-**90° and then flip them up-down to obtain the orientation corresponding to the SHWFS. Likewise, to use PD to validate a calibration acquired with the SHWFS we rotate the wavefront in the opposite direction (**+**90°) before flipping up-down to obtain the orientation corresponding to the EM-CCD.

To assess errors across multiple calibrations (**Fig. 3i**) we acquired three DM calibrations, each using a single bead located in a different field of view. For each acquired calibration (for both SHWFS- and PD-derived calibrations) we performed validation experiments using both sensing methods (**Supplementary Fig. 6**) across three different fields of view, resulting in a total of 9 validations for each sensing method. Each validation consists of a 20×20 matrix of coefficients, yielding 400 points per validation for a total of N = 3600 points each. Outliers were removed from validations using the generalized extreme Studentized deviate test for outliers with the MATLAB function `rmoutliers`, leaving a total of *N* = 3543 for the SHWFS and *N* = 3581 for the PD-derived calibration.

### Root mean square wavefront distortion

Wavefront aberrations are stated in terms of root mean square wavefront distortion of phase computed across *N* pupil elements relative to a flat wavefront according to

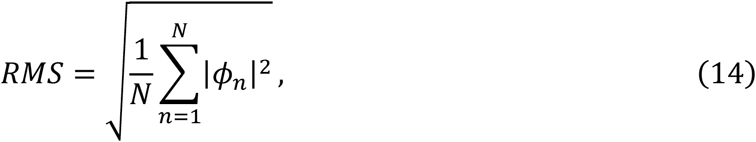

where *ϕ*_n_ is the phase for a given pupil position and *n* is the pupil pixel index.

### Random Aberration Generation and Scaling

When choosing aberrations for testing AO correction, Zernike coefficients were randomly generated mode-by-mode, with a 30% chance for the selected mode to have a non-zero value. Tip, tilt, defocus, and modes > 20 were excluded from selection. If selected, the random coefficient value *R*_*C*_ (in μm), was determined based on a variable amplitude, *R*_*A*_ via a random number *R*_*N*_ between 0 and 1, according to

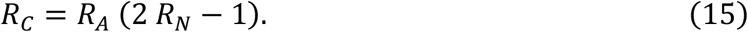

Values of R_C_ thus ranged from ±*R*_*A*_. We varied *R*_*A*_ between 0.1 to 0.8 when testing the effects of increasing aberration magnitude (**Fig. 4**); *R*_*A*_ was fixed to 0.4 when applying random aberrations for cellular imaging tests (**Fig. 5**).

In certain test cases, it was desirable to scale the magnitude of an aberration. The scaled coefficients, *C*_*sc*_ were obtained by multiplying each coefficient value *C*_*Z*_ by the ratio of the desired scaled wavefront RMS value, *RMS*_*scaled*_ and the original wavefront RMS value, *RMS*_*base*_:

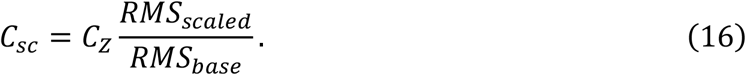

### AO Control Loop Software

AO correction was performed using a control loop implemented in the LabVIEW software program described above. A ground truth image (i.e., no aberration and system aberrations corrected) was acquired prior to starting the loop procedure. The AO correction loop was initialized by first generating and applying a random test aberration. The random aberration Zernike coefficients are then added to the diversity coefficients of each of the 5 total diversity phases. The four non-zero diversity phases consisted of pure horizontal or oblique astigmatism at ±1 μm amplitude (**Supplementary Fig. 7b**). Summed coefficients were converted to DM voltages using the DM control matrix and added to the offset voltages. The offset voltages served as the initial position for the deformable mirror, and were determined by removing system aberrations, described below.

After applying the relevant voltages to the DM, we acquired an image corresponding to each diversity phase. The laser was shuttered after acquisition of all diversity phases completed to prevent excess photobleaching, and the image stack and diversity phase information was passed to MATLAB for estimating the wavefront. After estimation, MATLAB passed the object estimate and estimated coefficients back to LabVIEW. These coefficients were converted to deformable mirror voltages using the control matrix and subtracted from the voltage previously applied for each phase, thereby applying AO correction. This process of acquisition, computation, and correction comprises one correction cycle. The process was repeated for a desired number of correction cycles.

### System Aberration Correction

The control loop described above can also be used to correct system aberrations (**Fig. 3j-m**). The primary difference in procedure is that for system aberration correction, no test aberrations are applied. Rather, an image is acquired, and the resulting wavefront sensed. Since this wavefront contains only system aberrations, we applied multiple correction cycles until the sensed wavefront error did not improve upon successive iterations. The voltages corresponding to the minimized wavefront error were saved and applied as the base voltage offset.

Axial views of 500 nm beads were acquired by applying the offset voltage to be tested (before or after correcting system aberrations) and acquiring a defocus image stack consisting of 101 image planes spaced 100 nm apart (**Fig. 3k-m**). Axial profiles were obtained using a binary mask to segment beads in the image area and computing the axial FWHM at the centroid of each segmented bead using the raw maximum intensity as the center position, using code available at https://github.com/eexuesong/RL_decon_spatial_domain/tree/main/FWHM.

### Testing Aberration Correction on Extended Samples

To test the capability of PD to correct aberrations induced in extended samples consisting of multiple 500 nm beads (**Fig. 4b-h**), the AO correction loop described above was applied n = 105 times with increasing amplitudes of randomly generated test aberrations for three, 512×512 fields of view selected to include at least 30 beads for a total of n = 315 tests. The selected aberrations were randomly generated in each test with the random amplitude *R*_*A*_ stepped from 0.1 to 0.8. We performed 10 tests each for *R*_*A*_ values from 0.1 to 0.3, and 15 tests for values from 0.4 to 0.8 to account for increased variability in the outcome of random coefficient values *R*_*C*_ at higher values of *R*_*A*_. For display purposes in **Fig. 4d, e** and to account for the random nature of the test aberrations, we binned the test aberration wavefront error in fixed 50 nm intervals. Note that x values displayed in **Fig. 4d, e** correspond to the mean values of the test aberration in each bin.

### Flat-Field Correction

To correct for non-uniform illumination, a flat-field correction was applied to cellular images (**Fig. 5**) after image acquisition. Flat-fielding was accomplished by dividing the original images by the average of 101 images of an autofluorescent plastic slide (92001, Chroma) and rescaling the quotient to a 14-bit value.

### Simulations based on synthetic images

To evaluate the performance of our PD method, we also conducted simulations with synthetic images (**Fig. 2a-c, Supplementary Figs. 2, 3**). We used three synthetic objects: 1) a resolution target (used for all simulations except those shown in **Fig 2b, c**); 2) images of microtubules acquired in a real microscope^44^ (**Fig 2b**); and 3) the Fishing Boat image (**Fig 2c**, from https://sipi.usc.edu/database/). Based on the forward imaging model (**Supplementary Note 1**), we added synthetic aberrations to these objects, creating aberrated images; generated additional diversity images with known diversity phases, and then used the PD algorithm to estimate the original aberrations. We generated random aberrations according to Eq. 15 for the first 20 Zernike modes (excluding piston, tip and tilt). In some cases, we also rescaled the aberrations to the desired magnitudes based on Eq. 16. Diffraction-limited images were generated by setting input aberrations to zero. The image size was set to 256 × 256 pixels in all cases, and simulations were performed within MATLAB.

When mimicking the imaging process, we also added mixed Poisson and Gaussian noise, generating images with SNR of

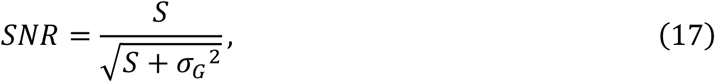

where S is the signal defined by the average of all pixels with intensity above a threshold (here set as 10% of the maximum intensity of the blurred objects in the noise-free image) and *σ*_*G*_ is the standard deviation of the Gaussian noise (here we used Gaussian noise with a mean value of zero and *σ*_*G*_ of 10). To determine the accuracy of wavefront estimation, we computed the root mean square (RMS) wavefront error of the difference between the PD-estimated wavefront and test aberrated wavefront (*ϕ*_*Residual*_).

In exploring the effect of parameter γ (in Eq. 1) on the performance of the PD algorithm (**Supplementary Fig. 2b**), we generated one raw aberrated image and one diversity image and used them to reconstruct the aberrated wavefront. The test aberrations were randomly generated and rescaled to have an RMS wavefront distortion of 150 nm. For a diversity phase, we used 0.5 μm amplitude astigmatism. The γ parameter was varied from 10^−1^ to 10^−30^, and for each γ setting, 50 independent simulations were carried out to obtain the mean and standard deviation of the RMS wavefront error. Simulations were repeated at 3 SNR levels (SNR = 5, 10 and 20).

To test different diversity phase choices (**Fig. 2d, Supplementary Fig. 3**), we generated one aberrated image and one diversity image, and used them to estimate the aberrated wavefront. Test aberrations were randomly generated and rescaled to an RMS wavefront distortion of 150 nm. The diversity phase was chosen from one of the following types of Zernike modes: defocus (Z=4), astigmatism (Z=5), coma (Z=8), trefoil (Z=9), spherical (Z=12), and a combination of defocus and astigmatism (Z=4 & Z=5). For each choice, we varied the RMS wavefront distortion of the diversity phase from 0.05 μm to 2 μm. For each diversity phase at each RMS value, 50 independent simulations (resolution target object, varying test aberrations and noise) were carried out to obtain the mean and standard deviation of the wavefront RMS errors (*ϕ*_*Residual*_). The SNR was set at 10 in these tests.

To investigate the effect of varying the number of diversity phases (**Fig. 2e**), we generated one aberrated image and different numbers of diversity images, using them to estimate the aberrated wavefront. The diversity phases were sequentially increased to a maximum of five: 1) astigmatism (Z=5) with amplitude of 0.5 μm; 2) astigmatism (Z=3) with amplitude of 0.5 μm; 3) astigmatism (Z=5) with amplitude of -0.5 μm; 4) astigmatism (Z=3) with amplitude of -0.5 μm; and 5) defocus (Z=4) with amplitude of 0.5 μm. In each simulation, the original test aberrations were randomly generated with RMS wavefront distortion varying from 0.02 μm to 0.24 μm. For each test aberration, 50 independent simulations were carried out to obtain the mean and standard deviation of the wavefront RMS errors (*ϕ*_*Residual*_). The SNR was set to 10 in these tests.

To explore the performance of the PD algorithm on images at different input noise levels (**Supplementary Fig. 2c**), we generated one aberrated image and four diversity images, and used them to estimate the aberrated wavefront. The four diversity phases were: 1) astigmatism (Z=5) with amplitude of 0.5 μm; 2) astigmatism (Z=3) with amplitude of 0.5 μm; 3) astigmatism (Z=5) with amplitude of -0.5 μm; and 4) astigmatism (Z=3) with amplitude of -0.5 μm. Test aberrations were randomly generated with RMS wavefront distortion varying from 0.02 μm to 0.24 μm. For each test aberration, 50 independent simulations were carried out to obtain the mean and standard deviation of the wavefront RMS errors (*ϕ*_*Residual*_). Simulations were conducted at SNR levels of 3, 5, 10, and 20 respectively.

## Competing Interests

C.J., M.G., P.J.L., and H.S. have filed a provisional patent for the technique described here.

## Author Contributions

Conceived project and directed research: H.S. Developed and implemented core phase diversity algorithm: M.G., N.R., Y.W., and P.J.L. Accelerated algorithm: C.J. and M.C.S. Wrote software: C.J., M.G., and M.C.S. Designed and built optical system: C.J., M.G., and H.S. Prepared samples: C.J., M.G., Y.S., S.K., and H.S. Performed experiments:. C.J. and M.G. Wrote paper: C.J., M.G., P.J.L, and H.S. with input from all authors. Supervised research: M.G., P.J.L., and H.S.

## Acknowledgements

This research was supported by the intramural research programs of the National Institute of Biomedical Imaging and Bioengineering, National Institutes of Health. We thank the Marine Biological Laboratories (MBL), for providing a meeting and brainstorming platform. H. S. and P. L.R. acknowledge the Whitman and Fellows program at MBL for providing funding and space for discussions valuable to this work. This work was supported by the Howard Hughes Medical Institute, and the Janelia Visiting Scientists Program. This research is funded in part by the Gordon and Betty Moore Foundation. This article is subject to HHMI’s Open Access to Publications policy. HHMI lab heads have previously granted a nonexclusive CC BY 4.0 license to the public and a sublicensable license to HHMI in their research articles. Pursuant to those licenses, the author-accepted manuscript of this article can be made freely available under a CC BY 4.0 license immediately upon publication. We thank Yanting Deng for initial coding efforts in LabVIEW; Xuesong Li for advice on optical alignment, bead sample preparation, and his comments on the manuscript; Cedric Allier for useful discussions about extending the work; and the trans-NIH Advanced Imaging and Microscopy (AIM) resource for housing the original prototype microscope constructed at NIH.

## Notes

### Summary of Updates

Fixed issue with figs clipping masks visible

