## Supplementary Information for "Phase diversity-based wavefront sensing for fluorescence microscopy"

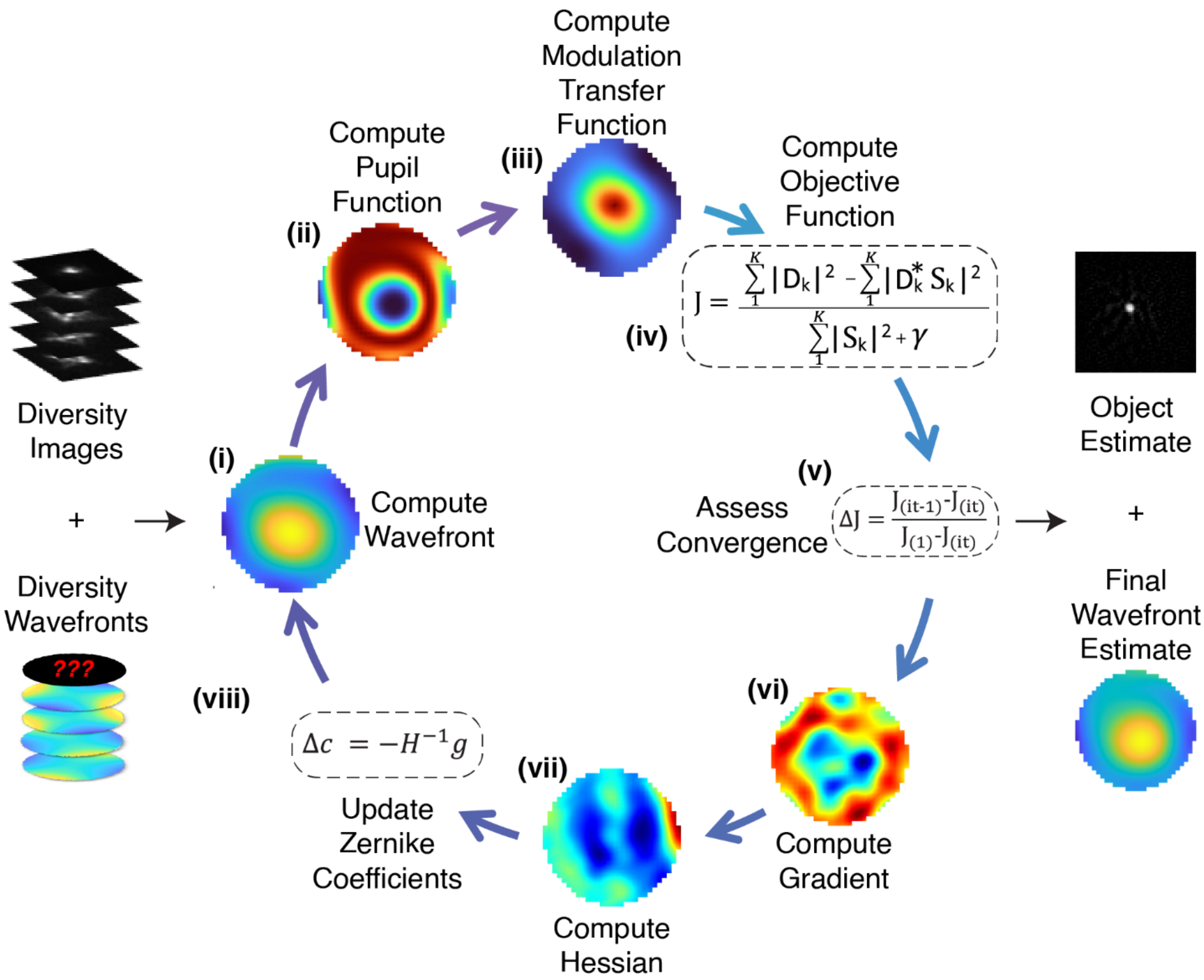


**Supplementary Fig. 1, Iterative wavefront estimation using phase diversity**. Process for iteratively estimating wavefront and object using input diversity images and corresponding diversity wavefronts. **(i)** Each loop starts by setting the wavefront to the previous iteration's wavefront estimate (in the first iteration the wavefront is set to zero), then computing the **(ii)** pupil functions and **(iii)** modulation transfer functions (MTF) associated with each of the diversity images, and ultimately the **(iv)** objective function to be minimized. If a **(v)** convergence criterion is met, wavefront and object estimates are returned. Otherwise, iteration continues by computing the **(vi)** gradient and **(vii)** Hessian, which are used to **(viii)** update the estimated Zernike coefficients, thereby estimating the current wavefront. See also **Methods**.

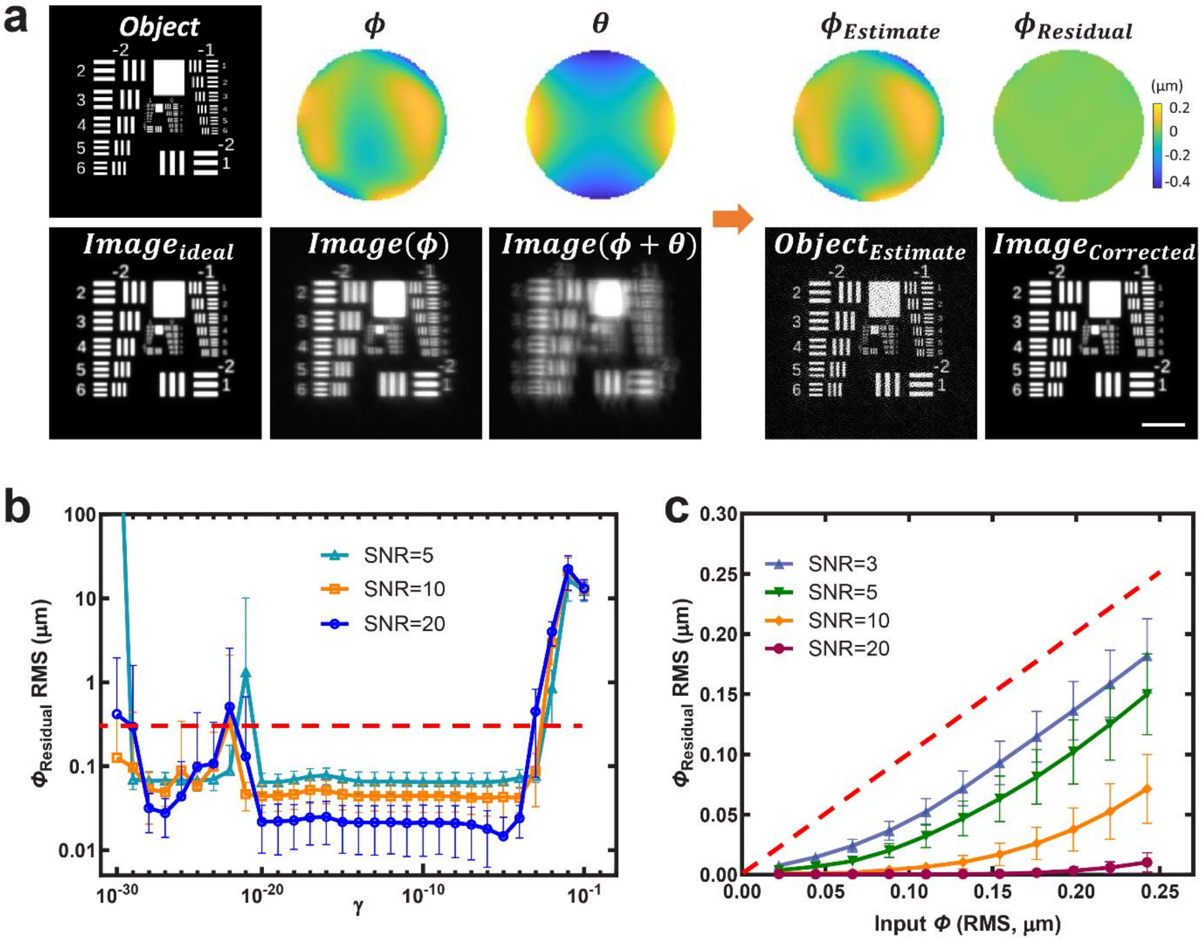


**Supplementary Fig. 2, Further simulations in support of our phase diversity algorithm. a)** Simulated adaptive optics using phase diversity. Left: when aberrating an object with wavefront $\phi$, an aberrated image *Image(*$\phi$*)* is produced instead of the ideal image $Image_{ideal}$. Acquiring an additional diversity image *Image(*$\phi+\theta$*)* with known diversity phase $\theta$ and using the phase diversity approach enables us to estimate the unknown phase $\phi_{Estimate}$ and the unknown object $Object_{Estimate}$. Subtracting $\phi_{Estimate}$ from $\phi$ yields the near-flat wavefront $\phi_{Residual}$ and generating the image associated with this residual phase simulates the process of AO, yielding $Image_{Corrected}$, which closely resembles $Image_{ideal}$. In this example, two input images are used in phase diversity (i.e., *Image(*$\phi$*)* and *Image(*$\phi+\theta$*)).* **b)** Investigating the role of the parameter $\gamma$ (see also **Methods**). We varied $\gamma$ to determine which range of values produced the best estimate of $\phi$, i.e., the lowest root mean square residual phase. Shown are simulation results for SNR levels of 5, 10, and 20. Means and standard deviations are shown for 50 simulations, each using two images (the original aberrated image and one diversity image with known diversity phase). Red dashed line indicates magnitude of input aberration (150 nm RMS wavefront distortion). **c)** Residual RMS wavefront error for different SNR levels, as a function of the magnitude of input aberration (see also **Fig. 2e**). Five images were used in these simulations (original aberrated image and four diversity images). Means and standard deviations from 50 simulations are shown, red dashed line shows y = x line;

values beneath this line correspond to cases in which the residual wavefront distortion is less than the input wavefront distortion. Scale bar in **a)**: 5 μm.

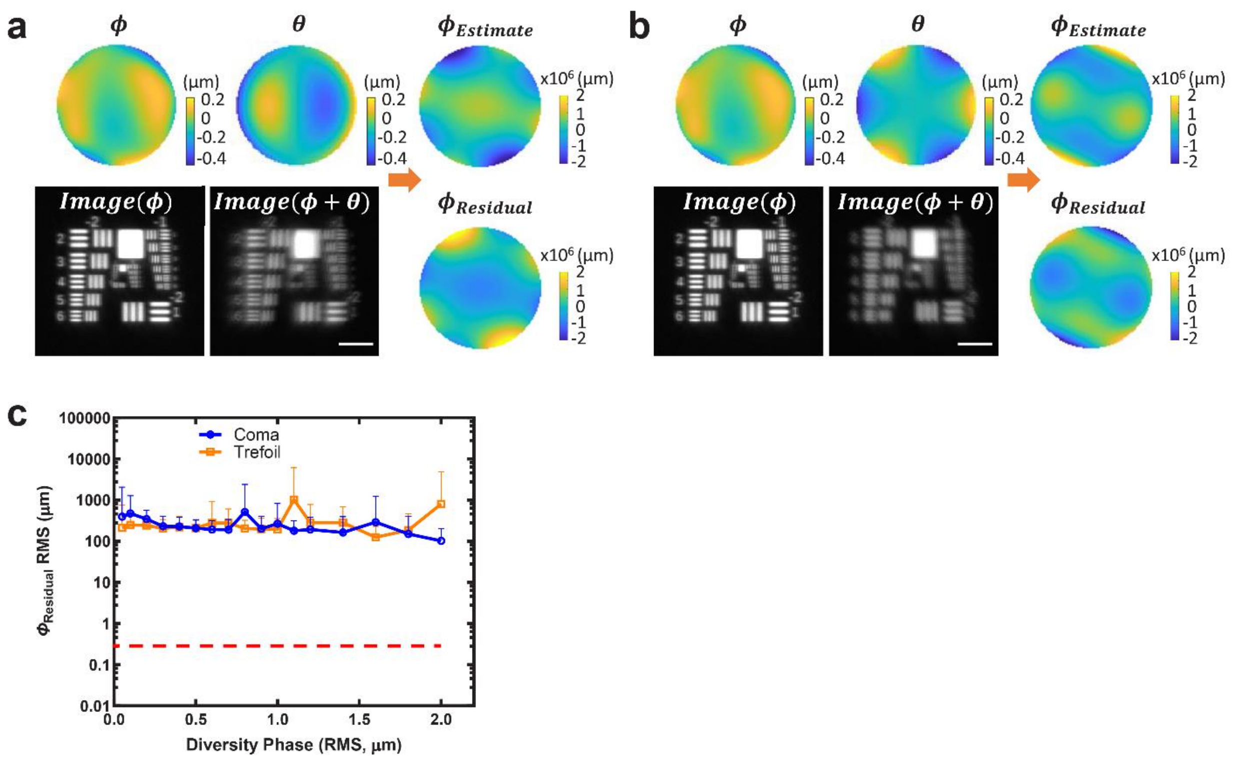


**Supplementary Figure 3, Coma and trefoil are poor diversity phase choices. a)** Aberration $\phi$, corresponding image *Image(*$\phi$), diversity phase $\theta$, and corresponding diversity phase *Image(*$\phi+\theta$) are used in phase diversity based wavefront sensing to produce estimate $\phi_{Estimate}$ with corresponding residual phase $\phi_{Residual}$. Here the diversity phase is coma. **b)** As in **a)** but using trefoil diversity phase. **c)** Both coma and trefoil produce estimated $\phi_{Residual}$ with RMS wavefront distortion far higher than the input aberration (red dashed line, note log scale on y axis). Results are shown as a function of RMS magnitude of the diversity phase, for 50 independent simulations. Scale bars in **a, b)**: 5 μm. Note that the different colorbars are used for different phase plots in **a, b**); the colorbar range for $\phi_{Residual}$ and $\phi_{Estimate}$ is orders of magnitude larger than that used for $\phi$ .

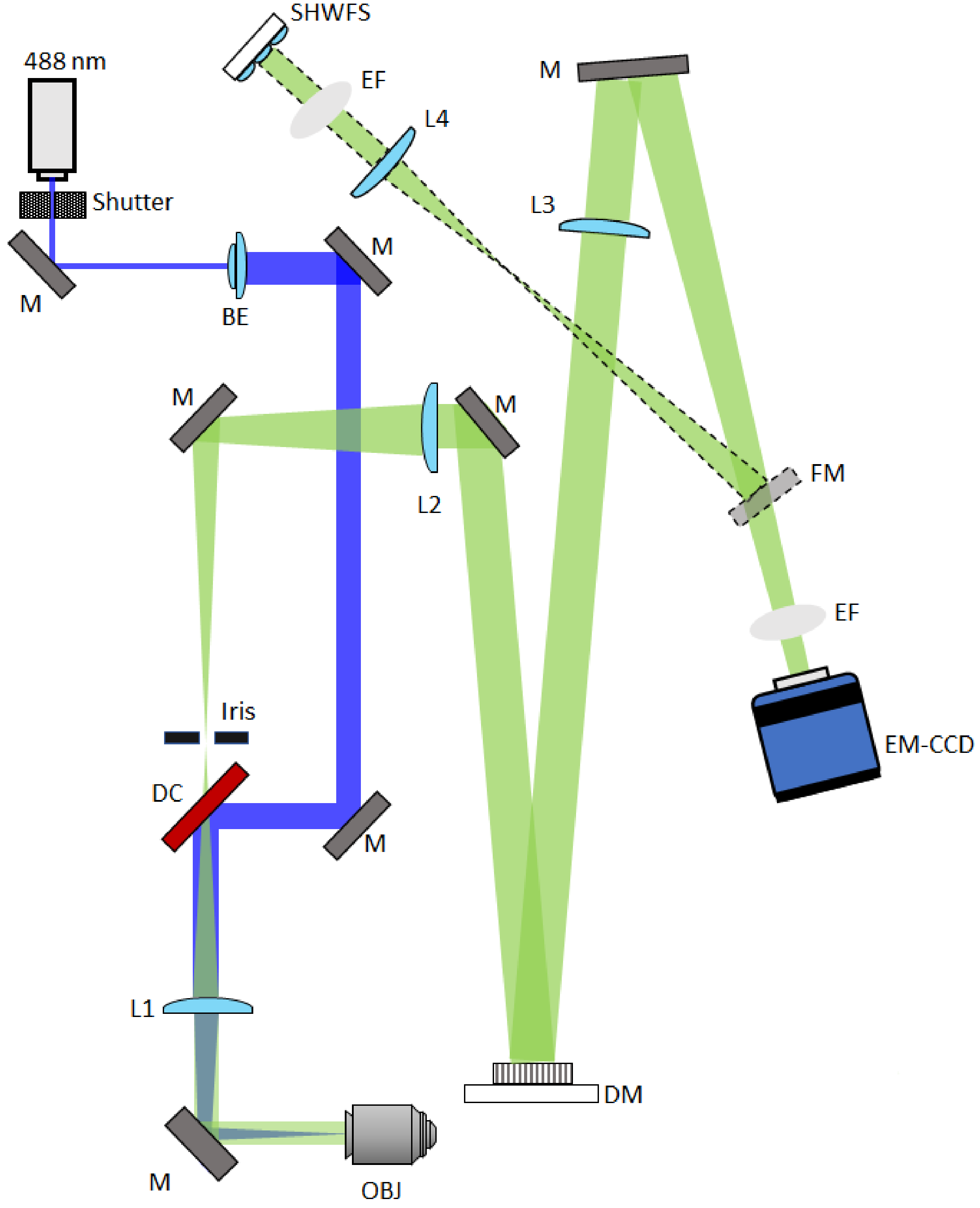


**Supplementary Fig. 4, Widefield adaptive optics setup used to test phase diversity algorithm.** The beam from a 488 nm laser (blue) was expanded 15x with a beam expander (BE), reflected from a dichromatic mirror (DC) and focused at the back focal plane of a 60x, 1.2 NA water objective (OBJ) with lens L1, thereby illuminating fluorescently labeled samples. Fluorescence (green) was collected in epi-mode through OBJ and L1 and separated from the illumination path via transmission through DC. A telescope consisting of lens L2 and L1 placed in 4f configuration served to magnify and image the objective pupil onto a deformable mirror (DM). Lens L3, placed one focal length after the DM then imaged the fluorescence onto an electron multiplying charge coupled device (EM-CCD), with fluorescence further isolated from illumination light via an emission filter (EF). Shuttering was achieved with a mechanical shutter and power attenuation via neutral density filters (not shown). In some experiments, the fluorescence was diverted (dashed green) via a flip mirror (FM), demagnified, and imaged to a Shack-Hartmann wavefront sensor (SHWFS) via lens L4, placed in 4f configuration to lens L3. In these experiments, we also placed an iris at an intermediate image plane upstream of the DM, to isolate the signal from a single fluorescence bead. Note that the figure is not to scale, but the number of reflections as indicated with turning mirrors (M) are accurate, and the layout of this figure approximates the setup used in this work. See **Methods** for additional information.

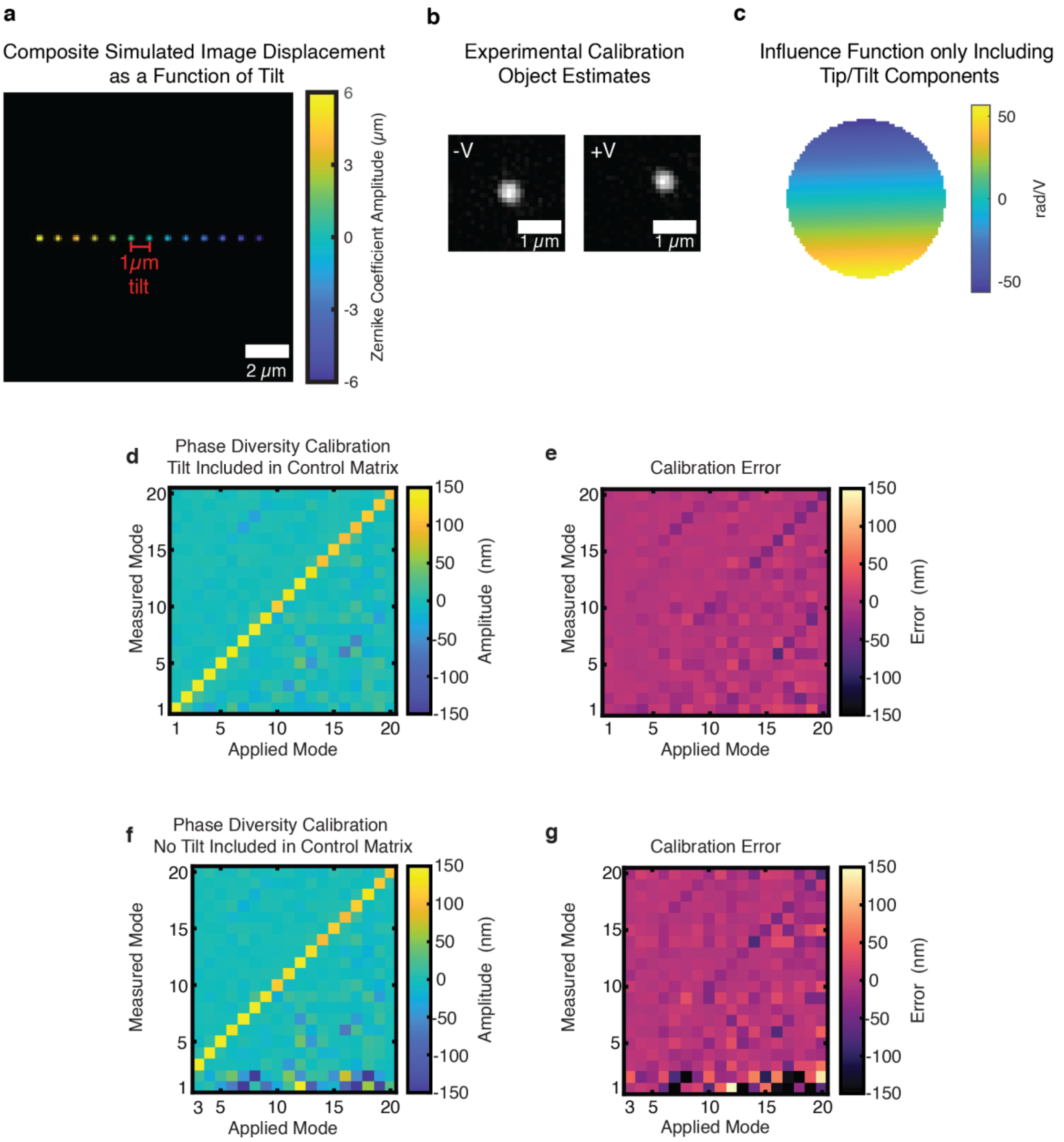


**Supplementary Fig. 5, Computing tip/tilt modes. a)** Simulation of object estimate displacement as a function of tilt yields the conversion factor δ, the amount the object moves in image space per unit amplitude tip/tilt coefficient. Colorbar representing amplitude of tip/tilt coefficient shown at right. **b)** Experimental object estimates at ±0.03 V show displacement due to residual tilt. **c)** Example calibration influence function showing tip/tilt components obtained from estimated object displacement. See also **Methods**. **d)** Characterization assay for phase diversity derived calibration measured using Shack-Hartmann wavefront sensor, with tip/tilt incorporated in control matrix. See also **Fig. 3h**. **e)** Error between applied and measured values for each entry in the characterization assay matrix in **d). f)** As in **d),** but without incorporating tip/tilt in the control matrix. Note the large values in the lowest two rows of characterization assay. **g)** Error for entries in the characterization assay matrix in **f)**.

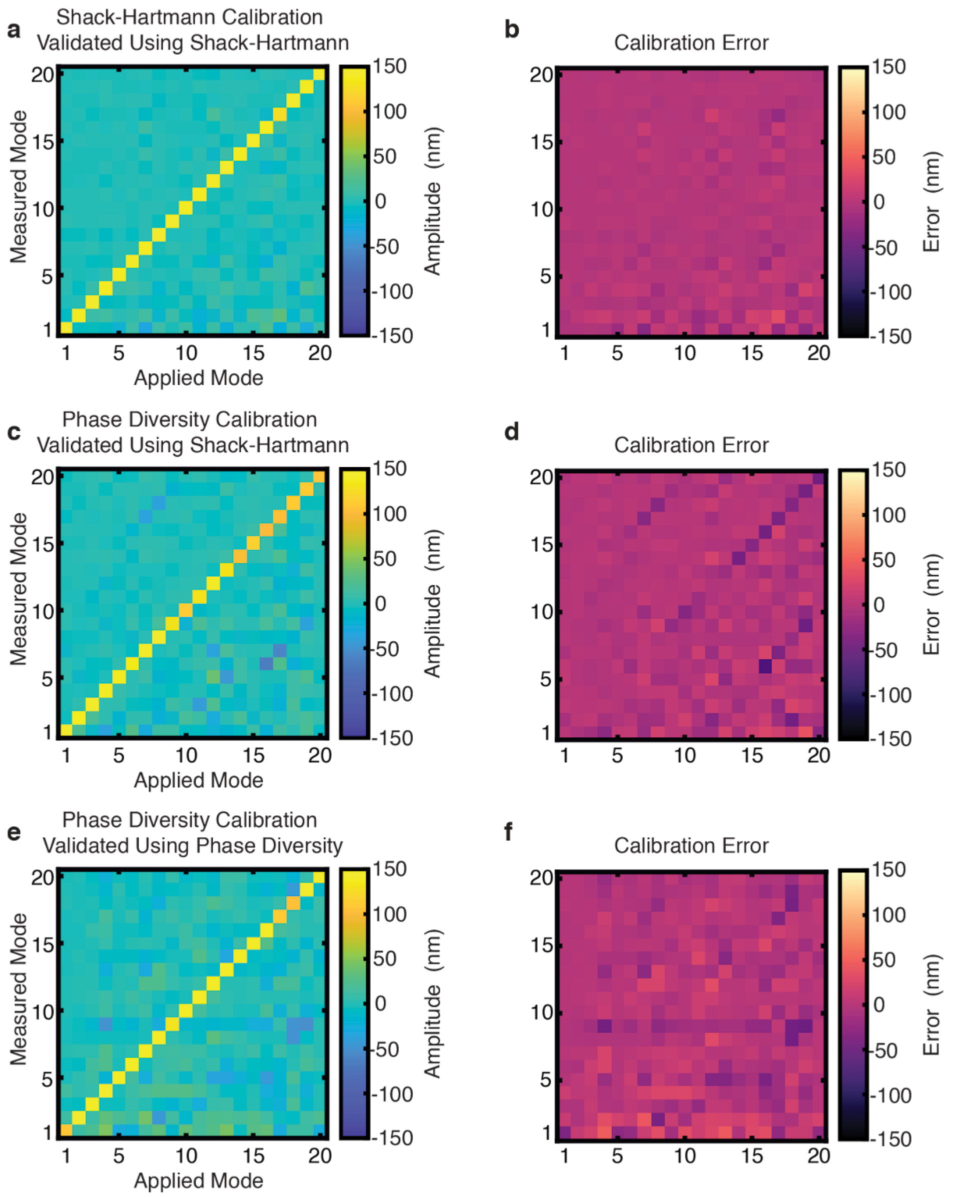


**Supplementary Fig. 6, Additional calibration data to accompany Fig. 3. a**) Replication of **Fig. 3g** with **b)** associated calibration error (i.e., difference) between expected and measured values for each entry in the characterization assay matrix. **c)** Replication of **Fig. 3h** with **d**) associated error. **e)** Example characterization assay for phase diversity derived calibration, measured using phase diversity. **f)** Error associated with **e)**. Note that values in error matrices **b, d**) are used to create violin plots shown in **Fig. 3i**).

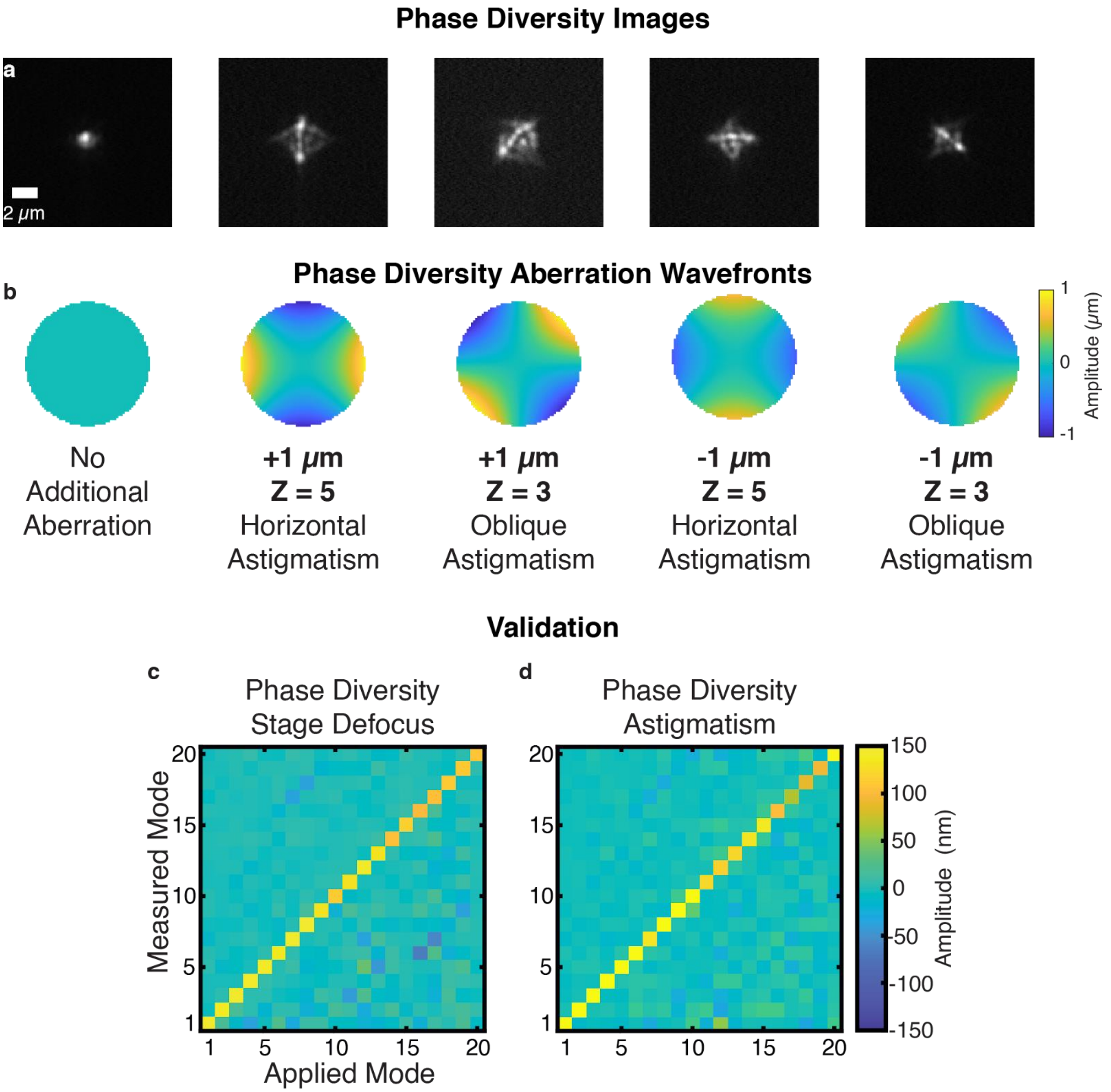


**Supplementary Fig. 7, Calibration using astigmatism diversity phases. a)** Phase diversity images using mirror-derived astigmatism aberrations (shown for a single voltage) and **b)** corresponding diversity wavefronts. **c)** Replicate of **Fig. 3h**) showing characterization assay of stage-defocus based DM calibration, versus **d)** calibration using astigmatism diversities. Off-target noise in the latter is reduced compared to the former.

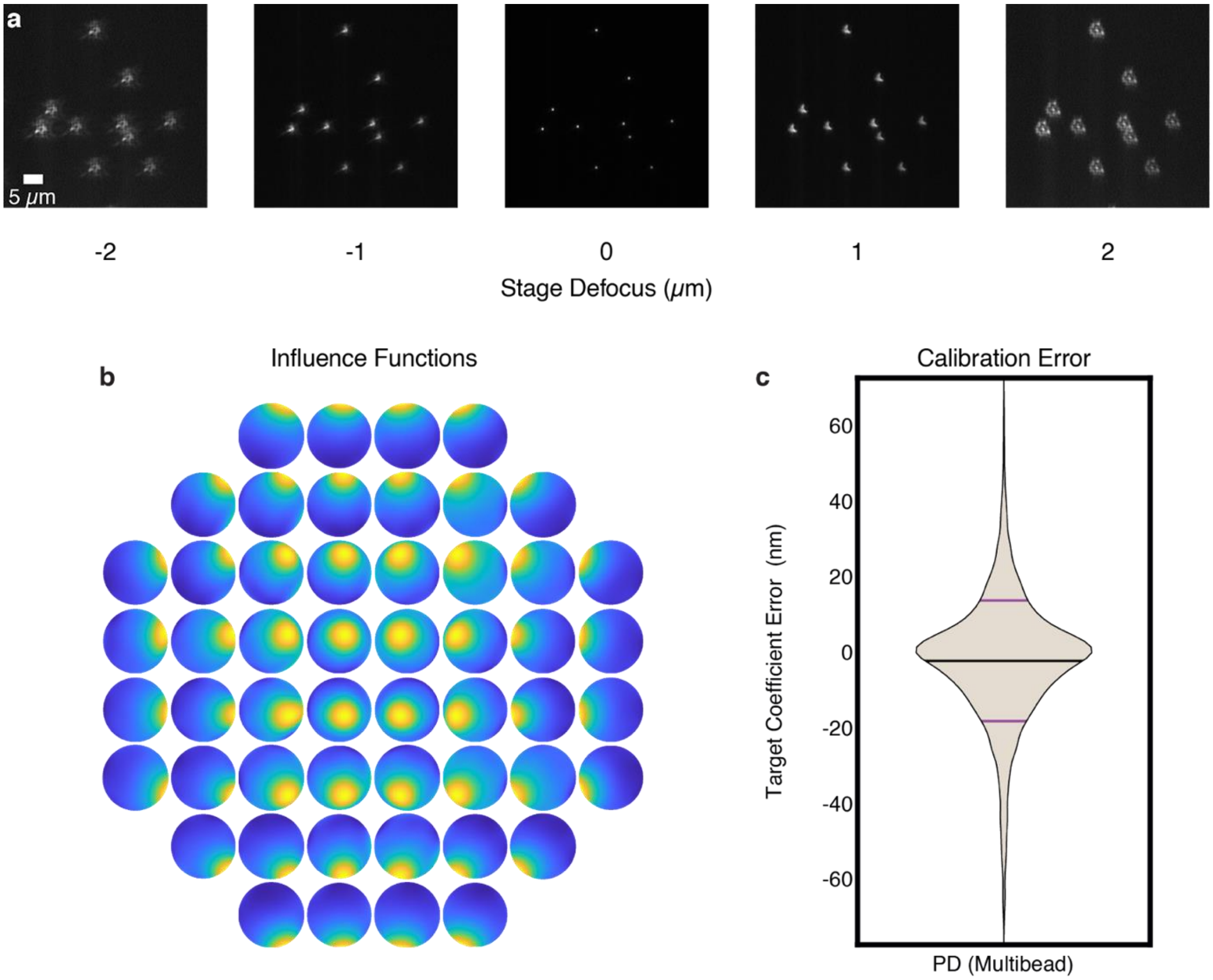


**Supplementary Fig. 8, DM calibration with multiple objects. a)** Example phase diversity image stack (single voltage shown) taken with multiple fluorescent beads. **b)** Influence functions derived from multi-bead stacks. **c)** Violin plot showing difference between target and measured coefficient values for each element in characterization matrix (not shown, measured with phase diversity). Mean (horizontal black bars) and standard deviations (horizontal red bars), -2.3 ± 16.0 nm. See also **Fig. 3i**.

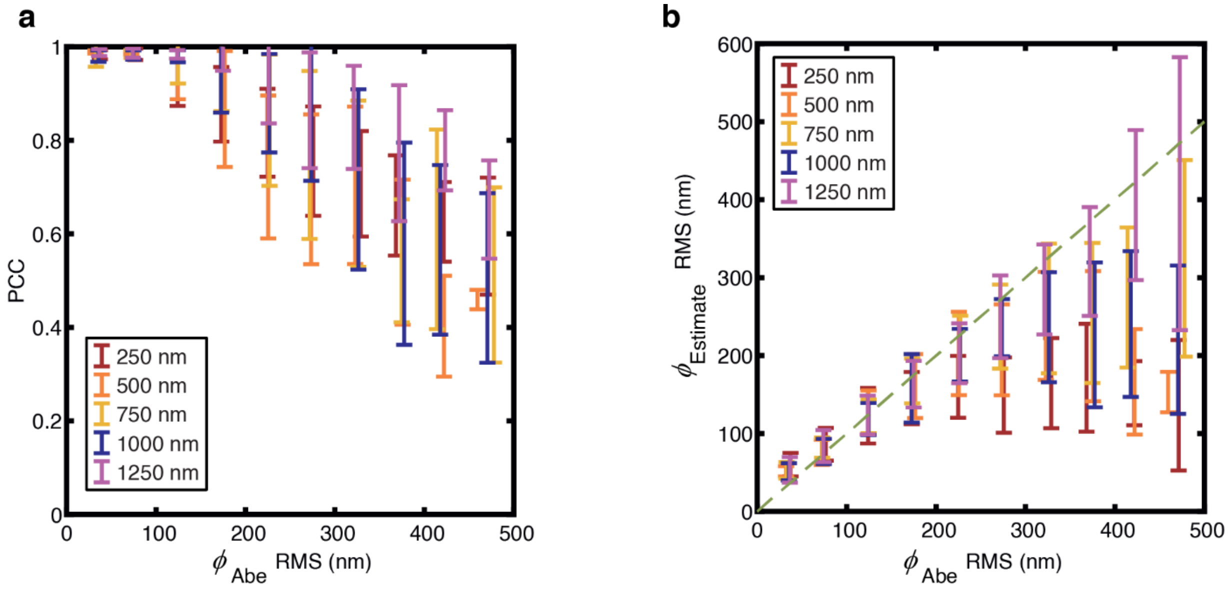


**Supplementary Fig. 9, Effect of different diversity magnitudes on correction and sensing.** Plots showing impact of magnitude of diversity aberrations (250 - 1250 nm) on corrected images **a)** and wavefront sensing **b).** As in **Fig. 4d, e)**, a random test aberration ($\phi_{Abe}$) was applied and a single cycle of sensing and correction was performed. **a**) Pearson correlation coefficient (PCC) between corrected and original unaberrated images for different diversity amplitudes, as a function of $\phi_{Abe}$ RMS wavefront distortion. **b)** Estimation of aberrated wavefront prior to correction. Dashed green line indicates values at which the estimated and test wavefront RMS values are equal. Data is pooled from N = 315 trials across 3 total fields of view for each diversity magnitude. Mean and SD y-axis values are shown for trials binned in 50 nm $\phi_{Abe}$ increments. X-axis values are shown centered at the mean values within each bin and may vary slightly due to random generation of aberrations.

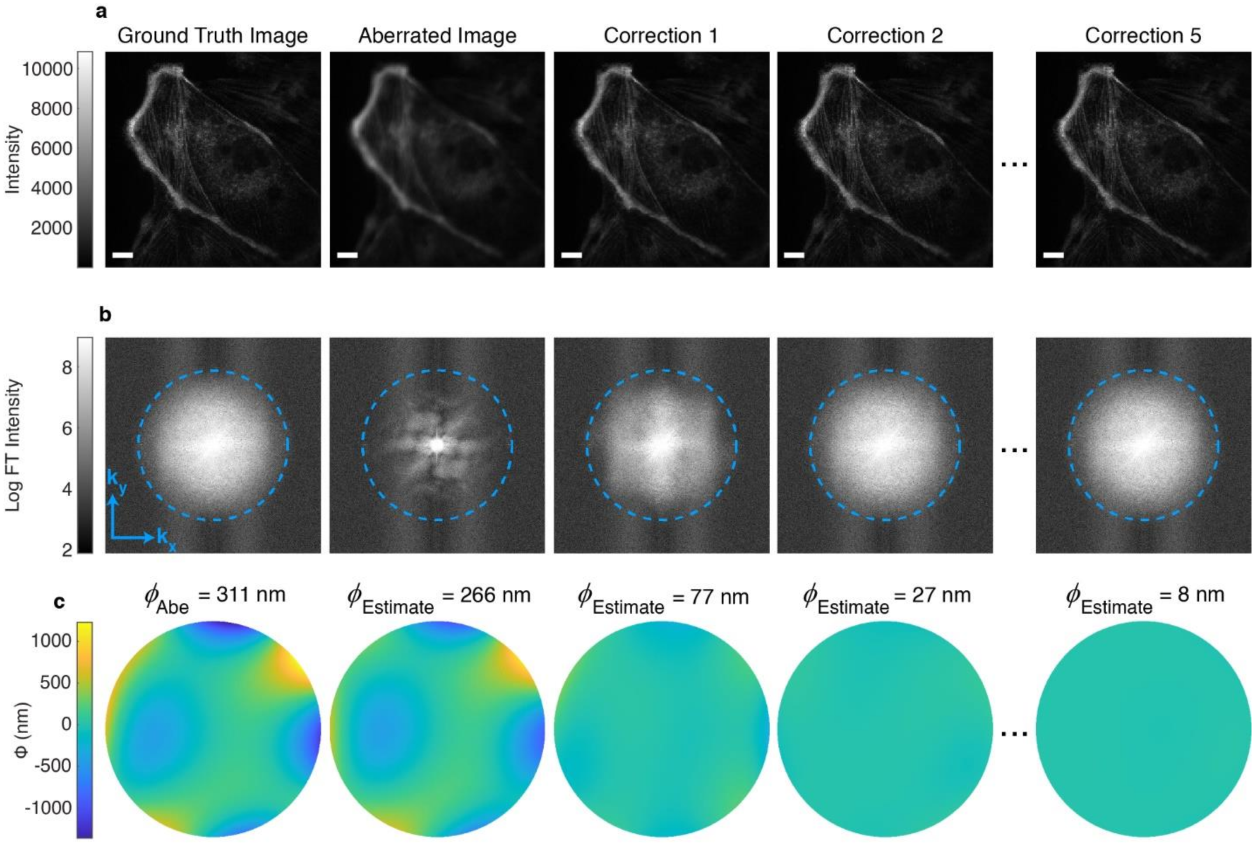


**Supplementary Fig. 10, Improvement in images of immunostained myosin as a function of correction cycle.** Image of fixed Ptk2 cells immunolabeled for myosin heavy chain, as in **Fig. 5i)**. Ground truth image, aberrated image, and AO-corrected image after first, second, and fifth correction cycle. Scale bar: 10 μm. **b)** Corresponding logarithm of magnitude of Fourier transform (FT). Dashed circles indicate 1/300 nm$^{-1}$. **c)** $\phi_{Abe}$ wavefront and $\phi_{Estimate}$ from phase diversity corresponding to aberrated and corrected images. RMS wavefront distortions are indicated above each wavefront.

**Supplementary Note 1: Phase diversity approach**

We pursue a computational approach to phase diversity that closely follows the mathematical strategy of Vogel et al.[1], though it differs in the parameterization of the problem. They focus on an astronomical situation in which aberrations are caused by temporal fluctuations in the atmosphere and they parameterize the phase error as a second-order, zero-mean stationary process. We instead represent the phase error in a microscopy system as an expansion in Zernike polynomials and seek to estimate the vector of coefficients of those polynomials. Because the use of a finite number of Zernike polynomials naturally leads to a reasonably smooth phase map, we omit the regularization term Vogel et al. apply to their phase map, but we do maintain the Tikohnov regularization term they apply to the estimated object.

We begin by establishing notation and the problem formulation. The pupil function $H_k(u)$ for the $k$th diversity image is given by

$$H_k(u) = |P(u)|e^{i\phi(u)+i\theta_k(u)},$$

where $u$ is a 2D pupil plane coordinate. Here $\phi(u)$ is the unknown phase aberration and $\theta_k(u)$, $k = 1\ldots, K$, is the $k$th known, purposefully introduced aberration. $|P(u)|$ is a binary pupil mask given by

$$P(u) = 0 \quad \text{for} \quad u > NA/\lambda,$$

where $NA$ is the numerical aperture of the objective and $\lambda$ the emission wavelength. The inverse Fourier transform of $H_k(u)$ is denoted by $h_k(x)$, where $x$ is the 2D spatial domain coordinate conjugate to $u$. For fluorescence imaging, the incoherent PSF will be given by

$$s_k(x) = |h_k(x)|^2.$$

The imaging model for the data acquired for the $k$th diversity image $d_k(x)$ is then given by

$$d_k(x) = s_k(x) * f(x) + n(x).$$

Here $f(x)$ is the unknown object of interest and $n(x)$ is additive noise.

The goal is to determine both $f(x)$, the unknown object, and $\phi(u)$, the unknown phase aberration. We assume that $\phi(u)$ can be expressed as the expansion of a finite number $M$ of Zernike polynomials $\phi_m(u)$ with coefficients $c_m$:

$$\phi(u) = \sum_{m=1}^{M} c_m \, \phi_m(u).$$

Thus, the aim is to determine the $M \times 1$ vector $\mathbf{c}$ of coefficients that best represents the phase map $\phi$, as well the unknown object $f(x)$.

**Iterative algorithm**

To find the phase map $\phi$ (by finding the vector of coefficients $\mathbf{c}$) and the object $f(x)$, we seek to minimize a Tikohonov-regularized least-squares objective function:

$$J = J[\phi, f] = \frac{1}{2}\left(\sum_{k=1}^{K} \int_{\mathbb{R}^2} [(s_k * f)(x) - d_k(x)]^2 \, dx\right) + \frac{\gamma}{2}\int_{\mathbb{R}^2} f\,(x)^2 dx.$$

It was shown by Gonsalves that there is a closed-form expression for the object (actually its Fourier transform) given an estimate of the phase map and the associated PSFs[2]. As given by Eq. 9 in Vogel,

$$F[\phi] = \frac{\sum_{k=1}^{K} S_k^*\,[\phi] D_k}{\gamma + \sum_{k=1}^{K} |S_k[\phi]|^2},$$

where $D_k$ are the Fourier transforms of the phase diversity images, $F[\phi]$ is the Fourier transform of the object $f(x)$ and

$$S_k[\phi] = \mathcal{F}\{s_k[\phi]\}$$

is the Fourier transform of the PSF. Note that for compactness we have dropped the explicit dependence on spatial position $x$ and spatial frequency $u$ but do preserve explicit dependence on $\phi$, which will be important to keep track of below when taking derivatives with respect to $\phi$. This allows the unknown object to be eliminated from the objective function, which can be reformulated in frequency space in terms of the unknown phase map only:

$$J = \frac{\sum_1^K |D_k|^2 \; - \; \sum_1^K |D_k^* S_k[\phi]|^2}{\sum_1^K |S_k[\phi]|^2 + \gamma}$$

To determine the phase map, Vogel et al. propose a Gauss-Newton method that is both fast and robust. It starts with an initial guess of the Zernike coefficients (typically $\mathbf{c} = \mathbf{0}$), with the update equation for the coefficients from iteration $q$ to $q + 1$ given by

$$\mathbf{c}^{(\mathrm{q}+1)} = \mathbf{c}^{(\mathrm{q})} + \Delta\mathbf{c},$$

where

$$\Delta\mathbf{c} = -\mathbf{H}^{-1}\mathbf{g}.$$

Here $\mathbf{g}$ is the $M \times 1$ gradient vector of the objective function and $\mathbf{H}$ is an $M \times M$ approximate Hessian (second derivative) matrix of the objective function.

**Gradient**

Vogel et al. show the expression for the gradient is given by

$$\mathbf{g}[\phi] = -2\sum_{k=1}^{K} \mathrm{Imag}\left\{H_k^*[\phi]\mathcal{F}\left\{h_k[\phi]\mathrm{Real}\left\{\mathcal{F}^{-1}\{V_k[\phi]\}\right\}\right\}\right\}.$$

In the above, $\mathrm{Imag}\{w\}$ denotes the imaginary part of a complex random variable $w$, $\mathrm{Real}\{w\}$ the real part, and $\mathcal{F}$ and $\mathcal{F}^{-1}$ denote the Fourier transform and its inverse, respectively. Also,

$$V_k[\phi] = F^*[\phi]D_k - |F[\phi]|^2 S_k[\phi].$$

The gradient has dimensions of the number of pixels in an image. To get the gradient with respect to the vector of coefficients $\mathbf{c}$, we simply need to take the inner product of the gradient above with the Zernike polynomials $\phi_m$:

$$g_m = \langle g[\phi], \phi_m \rangle, \qquad m = 1, \ldots M.$$

In summary, to calculate the gradient vector:

1. For $k = 1, \ldots K$, calculate the $D_k$ (FTs of the measured phase diversity images).
2. For the current estimate of $\mathbf{c}$, calculate $\phi = \sum_{m=1}^{M} c_m \, \phi_m$.
3. For $k = 1, \ldots K$, calculate $H_k[\phi] = |P| e^{i\phi + i\theta_k}$.
4. For $k = 1, \ldots K$, calculate $h_k[\phi]$ by taking the inverse FFT of $H_k[\phi]$.
5. For $k = 1, \ldots K$, calculate $s_k[\phi] = |h_k[\phi]|^2$.
6. For $k = 1, \ldots K$, calculate $S_k[\phi]$ by taking the FFT of $s_k[\phi]$.
7. Calculate $F$ using the closed-form expression above.
8. Calculate $V_k[\phi] = F^*[\phi]D_k - |F[\phi]|^2 S_k[\phi]$.
9. Calculate $\mathcal{F}^{-1}\{V_k[\phi]\}$ as needed for the gradient expression.
10. Evaluate the gradient expression.
11. Take the inner product with all $M$ Zernike polynomials to build the gradient vector $g$.

**The Hessian matrix**

To obtain the $M \times M$ Hessian matrix $\mathbf{H}$, we first adapt Eq. 26 in Vogel et al., which says that we can form the elements of $\mathbf{H}$ by acting with the operator $H$ on each of the $M$ Zernike polynomials and then taking inner products of the resulting $M$ vectors with each of the $M$ Zernike polynomials:

$$\mathbf{H}_{mn} = \langle H_{GN}[\phi]\phi_n, \phi_m \rangle.$$

Eq. 28 in Vogel et al. gives a way of evaluating $H_{GN}[\phi]\phi_n$:

$$H_{GN}[\phi]\phi_n = 4 \sum_{k=1}^{K} \sum_{j=1}^{k-1} \mathrm{Imag}\left\{ H_j^*[\phi] \mathcal{F}\left\{ h_j[\phi] \mathcal{F}^{-1}\{\widetilde{D}_k^* \, \tilde{U}_{jk}\}\right\} - H_k^*[\phi] \mathcal{F}\left\{ h_k[\phi] \mathcal{F}^{-1}\{\widetilde{D}_j^* \, \tilde{U}_{jk}\}\right\}\right\},$$

where

$$\tilde{U}_{jk} = \tilde{D}_j\,\mathcal{F}\left\{\mathrm{Imag}\left\{h_k^*[\phi]\mathcal{F}^{-1}\{H_k[\phi]\phi_n\}\right\}\right\} - \tilde{D}_k\,\mathcal{F}\left\{\mathrm{Imag}\left\{h_j^*[\phi]\mathcal{F}^{-1}\left\{H_j[\phi]\phi_n\right\}\right\}\right\},$$

with

$$\widetilde{D}_j = \frac{D_j}{Q^{1/2}}$$

and

$$Q[\phi] = \gamma + \sum_{k=1}^{K} |S_k[\phi]|^2.$$


1 Vogel, C. R., Chan, T. F. & Plemmons, R. J. Fast algorithms for phase-diversity-based blind deconvolution. *Proc. SPIE 3353, Adaptive Optical System Technologies* (1998).

2 Gonsalves, R. A. Phase retrieval and diversity in adaptive optics. *Optical Engineering* **21**, 829-832 (1982).
